# Reactivation of a dormant developmental plasticity program drives synaptic rejuvenation and antidepressant-like responses

**DOI:** 10.1101/2025.01.03.631260

**Authors:** Phi T. Nguyen, Emily Sun, Yuecheng Shi, Shogo Tamura, Yang Xiao, Clay Lacefield, Gergely F. Turi, René Hen

**Affiliations:** Departments of Psychiatry and Neuroscience, Columbia University Irving Medical Center; New York, NY 10032, USA; Division of Systems Neuroscience, New York State Psychiatric Institute; New York, NY 10032, USA; Department of Biomedical Engineering, Columbia University; New York, NY 10027, USA; Department of Systems Biology, Columbia University Irving Medical Center; New York, NY 10032, USA

## Abstract

Antidepressants are widely used to treat mood and anxiety disorders, yet the cellular and molecular mechanisms underlying their therapeutic effects remain poorly understood. Here, we show that the selective serotonin reuptake inhibitor (SSRI) fluoxetine rejuvenates the dentate gyrus (DG) by reactivating a latent developmental plasticity program in mature neurons, which is required for re-engaging critical period-like synaptic remodeling and antidepressant-like behavioral effects. Single nuclei transcriptomic profiling across all hippocampal cell types revealed a strikingly selective response in the DG, where mature neurons re-engaged a developmental-like transcriptional state, an effect distinct from adult neurogenesis. Chronic fluoxetine treatment induced expression of SOX11, a developmental transcription factor involved in neurodevelopment and axonal regeneration that is normally silenced in the adult brain. Reactivation of SOX11 in mature DG neurons was necessary for both the antidepressant-like behavioral effects of fluoxetine and the re-engagement of critical period-like synaptic remodeling. Notably, SOX11 reactivation was also induced by physiological stimuli, including environmental enrichment and peripheral axon regeneration, suggesting that mature neurons retain access to a conserved developmental plasticity program throughout adulthood. We further identified evidence of this form of developmental plasticity in the adult human brain, an effect distinct from adult neurogenesis. At the structural level, fluoxetine remodeled novel extracellular matrix (ECM) structures within the DG distinct from canonical perineuronal nets, and targeted ECM degradation was sufficient to reactivate SOX11 in mature neurons, identifying the ECM as a key regulator of neuronal rejuvenation. Altogether, these findings demonstrate that mature neurons retain an intrinsic capacity to re-engage dormant developmental programs, and that antidepressants harness this regenerative plasticity to drive neural circuit remodeling and antidepressant-like effects.

## Main

Mood and anxiety disorders, including major depressive disorder (MDD), are among the most prevalent mental illnesses and a leading cause of disability worldwide^1^. First line treatments for mood disorders include SSRIs such as fluoxetine, but despite decades of clinical use, they achieve remission in less than half of patients and usually require several weeks to achieve their full therapeutic effects^2^. This limited efficacy underscores the critical need to define the cellular and molecular mechanisms underlying the antidepressant response.

At the pharmacological level, SSRIs primarily act by inhibiting 5-HT reuptake and increasing serotonergic tone, yet this effect occurs within hours whereas the antidepressant response requires several weeks to manifest. This temporal effect has led to the hypothesis that therapeutic efficacy depends on slower forms of neural plasticity, including neurotrophin-dependent synaptic remodeling and increased adult neurogenesis^3–5^. Interestingly, SSRIs can also reinstate features of developmental and critical period plasticity in the adult brain, such as ocular dominance plasticity in the visual cortex^6–9^. However, the molecular programs that enable the reactivation of developmental plasticity in the adult brain remain poorly defined, and whether this reactivation is causally required for antidepressant effects on mood-related behavior is unknown. Defining these mechanisms may therefore reveal new therapeutic targets for enhancing antidepressant responses.

Emerging evidence implicates the extracellular matrix (ECM) as a regulator of latent plasticity states. During development, the formation of ECM structures such as perineuronal nets (PNNs) contributes to the closure of critical periods^10^, while ECM removal can restore features of critical period plasticity in the adult brain^11–14,6,7,15,8,16,9^. In the adult brain, chronic stress and antidepressant treatment bidirectionally remodel the ECM^17–20^. However, the underlying molecular programs and their relationship to the pathways mediating antidepressant responses remain poorly understood. This is, in part, because most studies have focused predominantly on PNNs, which represent only a small fraction of the total ECM, and on transcriptional measures that do not capture the long-lived nature of ECM proteins^21^. Defining the mechanisms underlying ECM-gated plasticity therefore holds significant promise for understanding the neurobiology of antidepressant responses and for guiding the development of targeted therapeutic strategies.

### Fluoxetine induces time-dependent transcriptomic reprogramming selectively in mature dentate granule cells

To investigate the cellular mechanisms by which SSRIs modify stress-related behaviors, we used a well-established mouse model of chronic stress and fluoxetine (Flx) treatment^22,23^ (**Fig. 1A**). In this paradigm, administration of corticosterone (Cort), the primary stress hormone in rodents, recapitulates behavioral impairments associated with chronic stress, a major risk factor for mood and anxiety disorders^24^. Mice received chronic Cort via drinking water for 6 weeks, with a subset receiving concurrent Flx treatment for either 3 weeks or 5 days. Consistent with previous studies^22^, Cort-treated mice in the elevated plus maze (EPM) showed reduced exploration of the anxiogenic open arm, an effect that was reversed by 3 weeks of Flx (**Fig. 1B** and **Fig. S1A**). In contrast, 5 days of Flx had no effect, mirroring the delayed onset of therapeutic effects observed in patients. In the novelty-suppressed feeding (NSF) test, an anxiety-related conflict test, Cort-treated mice showed an increased latency to feed in the anxiogenic arena, which was rescued by 3 weeks, but not 5 days, of Flx treatment (**Fig. 1C** and **Fig. S1B**). Three weeks of Flx treatment also elicited antidepressant-like behavioral responses in the forced swim test (**Fig. S1C**), consistent with previous studies^22^. These results therefore provide a foundation to investigate the molecular programs underlying the effects of Flx.

**Fig. 1.**
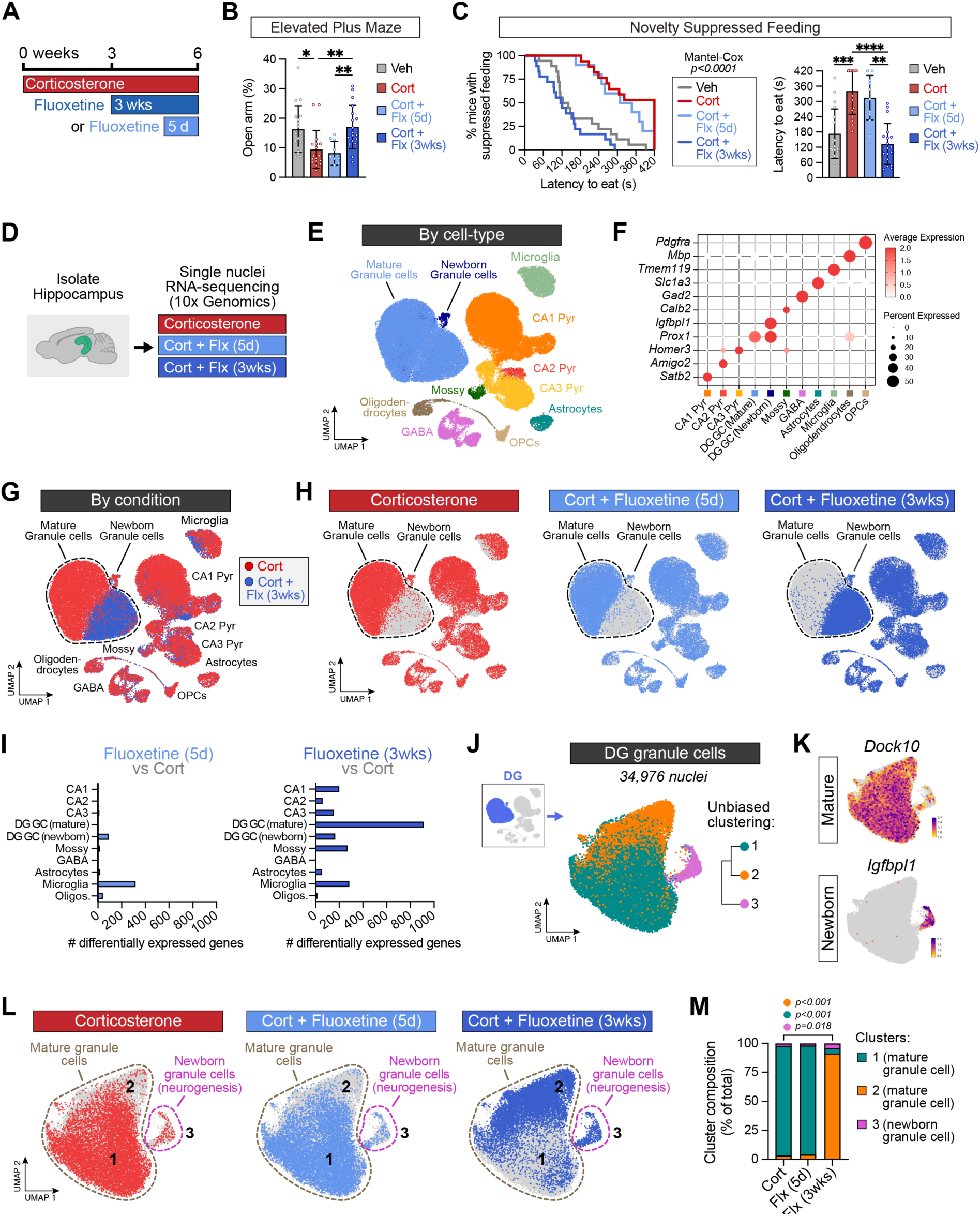
Fluoxetine induces transcriptomic reprogramming selectively in mature neurons of the dentate gyrus. (**A**) Pharmacological approach. Corticosterone (35 ug/mL) and Fluoxetine (160 ug/mL) were delivered in drinking water. (**B**) Time spent in the open arm of the EPM per mouse (One-way ANOVA, p=0.0007, Tukey’s post-hoc test, n=18 (Veh), 17 (Cort), 10 (Flx 5d), 18 (Flx 3wks)). (**C**) Latency to feed in NSF per mouse shown as survival curve (left; Log-rank Mantel-Cox test, χ2=39.21, p<0.0001) and bar plot (right; Kruskal-Wallis ANOVA, p<0.0001, Dunn’s post-hoc test, n= same as (B)). (**D**) Paradigm for downstream snRNA-seq of hippocampal cells (n=3 mice per condition). (**E**) Unsupervised clustering (uniform manifold approximation and projection (UMAP)) of 119,346 single nuclei transcriptomic profiles labeled by cell type. (**F**) Expression of cell type-specific marker genes for major neuronal and glial populations. See Fig. S2D for additional marker genes. (**G**) Nuclei from Cort + Flx (3 wks) group (blue) overlaid with nuclei from Cort alone (red). (**H**) Nuclei highlighted by treatment condition. (**I**) Number of differentially expressed genes (DEGs) between Cort and Cort + Flx groups for each cell type. Each cell type was downsampled to 588 nuclei per mouse for all analyses. See also *Methods*. (**J**) Subclustering of 34,976 dentate granule cells from (E). (**K**) Expression of maturation-related marker genes in granule cells. (**L**) Granule cell nuclei highlighted by treatment condition. (**M**) Quantification of granule cell cluster composition by treatment condition (Two-way ANOVA, condition p>0.9999, cluster p<0.0001, interaction p<0.0001, Tukey’s post-hoc tests, n=3 mice per condition). Data = mean ± SD. \**p < 0.05*; \*\**p < 0.01*; \*\*\**p < 0.001*; \*\*\*\**p < 0.0001*.

The hippocampus is strongly implicated in the pathophysiology of stress-related disorders^24–27^ and is required for the behavioral effects of Flx in rodent models^5,22,28,29^. To define the molecular substrates in the hippocampus driving behavioral responses to Flx, we performed droplet-based single nucleus RNA-sequencing (snRNA-seq, 10x Genomics v3) of hippocampal nuclei in mice treated with Cort or Cort + Flx for 5 days or 3 weeks (**Fig. 1D**). We recovered 119,346 high-quality nuclei across all conditions and identified the major neuronal and glial subtypes in the hippocampus for downstream analyses (**Fig. 1E-F**, and **Fig. S2**).

Unsupervised clustering analysis revealed a striking transcriptomic shift between Cort and Cort + Flx (3wks) groups, which predominantly mapped onto DG granule cells (GCs) (**Fig. 1G-H**). Although the DG contains both mature GCs and newborn GCs generated through adult hippocampal neurogenesis, this shift occurred primarily in the mature population and was not present after 5 days of Flx. To assess the extent of transcriptomic reprogramming, we quantified the number of differentially expressed genes (DEGs) between Cort and Cort + Flx groups for each cell type (**Fig. 1I**). After 5 days of Flx, transcriptional changes were minimal across most cell types, with the exception of microglia, consistent with the lack of behavioral effects at this time point (**Fig. 1B-C**). In contrast, neuronal responses became dominant after 3 weeks of Flx, with mature GCs producing the largest number of DEGs, an effect that was at least ∼4-fold greater than any other neuronal population (**Fig. 1I**). Altogether, these results suggest that Flx induces time-dependent transcriptomic reprogramming selectively in mature DG GCs.

Both mature and newborn GCs are required for the behavioral effects of chronic SSRI treatment in rodents^5,22,28^. However, whether the underlying mechanisms diverge across these heterogenous populations remains poorly understood. To define the molecular responses of these distinct populations, we leveraged the single cell resolution of our dataset to subcluster GCs (34,976 nuclei), revealing three transcriptionally distinct clusters (**Fig. 1J**). Clusters 1 and 2 were identified as mature GCs, characterized by high *Dock10* and low *Igfbpl1* expression, whereas cluster 3 corresponded to *Igfbpl1*+ newborn GCs^30^ (**Fig. 1K**). In the Cort and Cort + Flx (5d) groups, mature GCs mapped primarily onto cluster 1, suggesting minimal transcriptional changes at this early timepoint (**Fig. 1L-M**). However, in the Cort + Flx (3 wks) group, mature GCs mapped primarily onto cluster 2, consistent with the emergence of a distinct cellular state. Newborn GCs remained transcriptionally stable across conditions, but expanded in number ∼4-fold after 3 weeks of Flx, consistent with previous findings^4^. Thus, while previous models of antidepressant effects have focused on newborn GCs generated by adult neurogenesis, our results indicate that among all hippocampal cell types, mature GCs undergo the most striking transcriptomic reprogramming in response to chronic Flx.

### Fluoxetine re-engages neurodevelopmental plasticity programs in mature neurons independent of hippocampal neurogenesis

Neurodevelopment is characterized by critical periods of heightened plasticity that promote large-scale neural circuit reorganization. Previous studies have shown that SSRIs can reactivate features of critical period plasticity in the adult brain, including ocular dominance plasticity in the visual cortex and enhanced axon regeneration in the spinal cord^6–8,15^. However, the underlying molecular mechanisms by which Flx exerts these critical period-like effects, and how these mechanisms relate to antidepressant efficacy, remain poorly understood.

To define the molecular programs by which mature GCs respond to SSRIs, we analyzed this population separately from newborn GCs and ranked DEGs between Cort and Cort + Flx (3 wks) groups (**Fig. 2A-B** and **Table S1**). Gene ontology analysis revealed significant enrichment among upregulated genes for pathways implicated with synaptogenesis and neurodevelopment (*Sox11*, *Bdnf*, and *Sema3e*) (**Fig. 2B-C**). Consistent with previous studies^9^, Flx also downregulated genes associated with neuronal maturity, such as *Calb1*, *Dock10*, and *Stxbp6*. These findings indicate that mature neurons in the DG can re-engage features of neurodevelopment independent of adult neurogenesis.

**Fig. 2.**
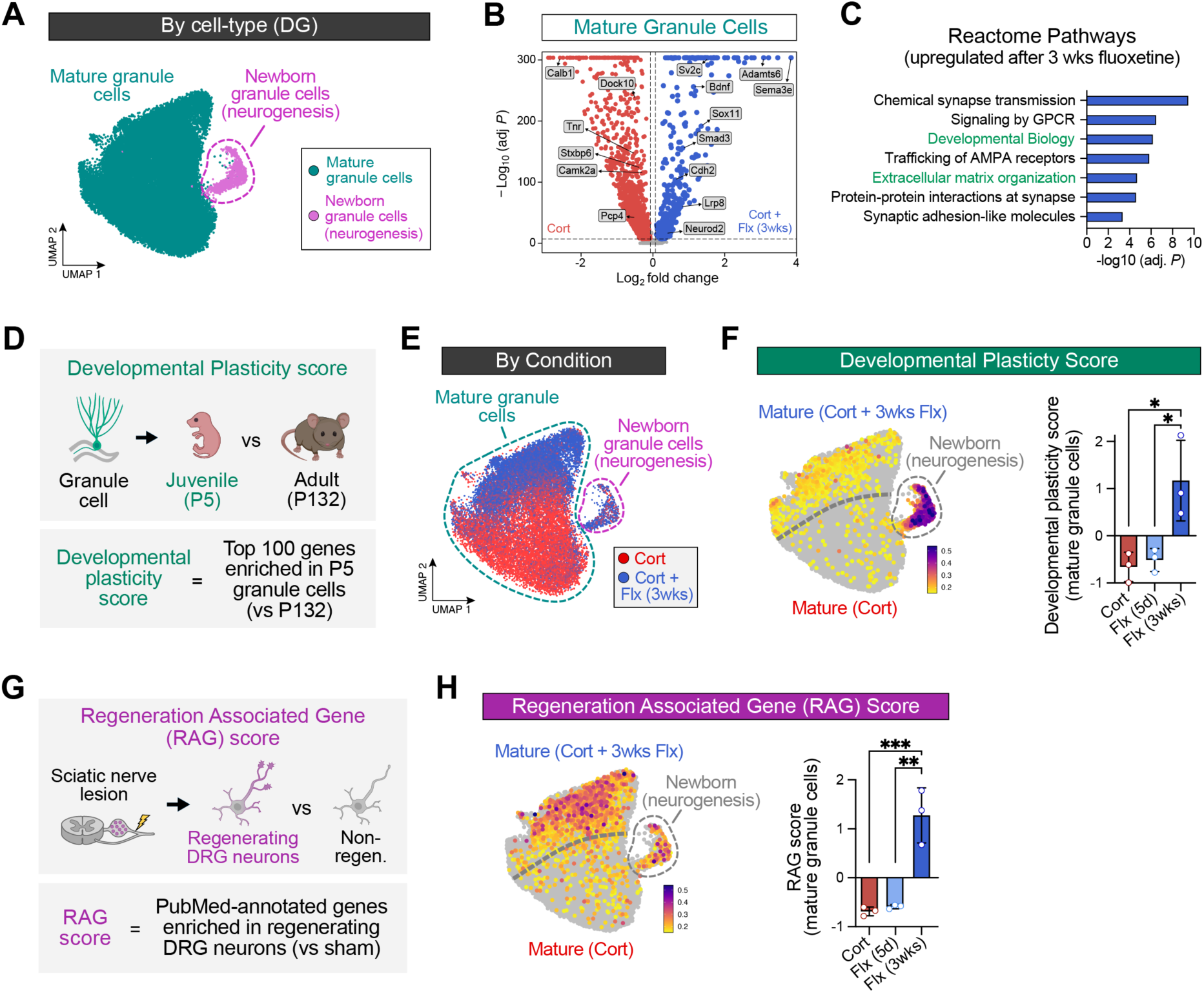
Fluoxetine reprograms mature neurons to a developmental-like state distinct from adult neurogenesis. (**A**) Identification of mature and newborn DG granule cell populations. (**B**) Volcano plot of differentially expressed genes selectively from mature granule cells from Cort and Cort + Flx (3wks) groups. (**C**) Top gene ontology terms for upregulated genes in (B). (**D**) Developmental Plasticity Score generated from the top 100 genes upregulated in P5 granule cells versus P132 granule cells. Dataset from *Hochgerner et al., 2018*^31^; see also Fig. S3A-B, Table S2, and *Methods*. (**E**) Transcriptomic profiles of dentate granule cells colored by treatment condition (Cort = red, Cort + Flx (3wks) = blue). (**F**) Feature plot showing average expression of “Developmental Plasticity Score” (left) and z-scores selectively in mature granule cells per mouse (right; One-way ANOVA, p=0.0110, Tukey’s post-hoc test, n=3 mice/group). (**G**) Regeneration Associated Gene Score generated from WGCNA-derived, PubMed-annotated consensus genes associated with regenerating dorsal root ganglion (DRG) neurons. Dataset from *Chandran et al., 2016*^39^; see also Fig. S3C, Table S2, and *Methods*. (**H**) Feature plot showing average expression of “Regeneration Associated Gene Score” (left) and z-scores selectively in mature granule cells per mouse (right; One-way ANOVA, p=0.0005, Tukey’s post-hoc test, n=3 mice/group). Data = mean ± SD. \**p < 0.05*; \*\**p < 0.01*; \*\*\**p < 0.001*

To test whether chronic Flx treatment re-engages neurodevelopmental plasticity programs in mature neurons, we analyzed a scRNA-seq dataset spanning DG development^31^ and identified DEGs between GCs from juvenile (P5) and adult (P132) mice (**Fig. 2D** and **Fig. S3A-B**). We then generated a “Developmental Plasticity Score” using the top 100 genes upregulated in P5 GCs (**Table S2**). As expected, newborn GCs generated through adult neurogenesis showed a high “Developmental Plasticity Score” (**Fig. 2E-F**), consistent with findings that adult neurogenesis recapitulates features of neurodevelopment^32^. Strikingly, mature GCs from Cort + Flx (3 wks) mice also showed a significant increase in the “Developmental Plasticity Score”. This effect was absent at 5 days of Flx, mirroring the delayed onset of SSRI-induced behavioral effects (**Fig. 1B-C**). These findings suggest that mature neurons in the adult brain retain an intrinsic capacity to re-engage latent developmental transcriptional programs.

Outside the brain, mature neurons in the PNS can re-enter a developmental-like state after nerve injury to facilitate axon regeneration^33–35^. This regenerative response depends on the induction of a transcriptional program comprised of regeneration associated genes (RAGs), including developmental transcription factors such as *Sox11*^36–38^. To assess whether SSRI-induced plasticity engages similar developmental-like mechanisms, we generated a “Regeneration Associated Gene Score” based on canonical RAGs^39^ (**Fig. 2G**, **Fig. S3C**, and **Table S2**). Following 3 weeks of Flx treatment, mature GCs showed a significant increase in the “Regeneration Associated Gene Score” (**Fig. 2H**), suggesting that Flx induces a regeneration-like transcriptional state in mature neurons that supports axonogenesis and structural plasticity. Altogether, our results indicate that chronic Flx drives mature GCs to reactivate a dormant developmental plasticity program, providing a candidate molecular substrate for the antidepressant-like effects of SSRIs.

### Sox11 is a conserved developmental switch that is reactivated in mature neurons during physiological contexts and diverse antidepressant interventions

We next investigated the molecular drivers that reactivate this dormant developmental plasticity program in mature GCs following chronic Flx treatment. Because transcription factors are well positioned to coordinate broad transcriptional networks, we used transcription factor enrichment analysis to identify factors whose DNA binding motifs were overrepresented among genes induced by chronic Flx (**Fig. 3A**). This analysis revealed SOX11 among the top transcription factors predicted to regulate Flx-induced genes. Accordingly, we found that *Sox11* expression was significantly increased in mature GCs after 3 weeks of Flx treatment (**Fig. 3B**).

**Fig. 3.**
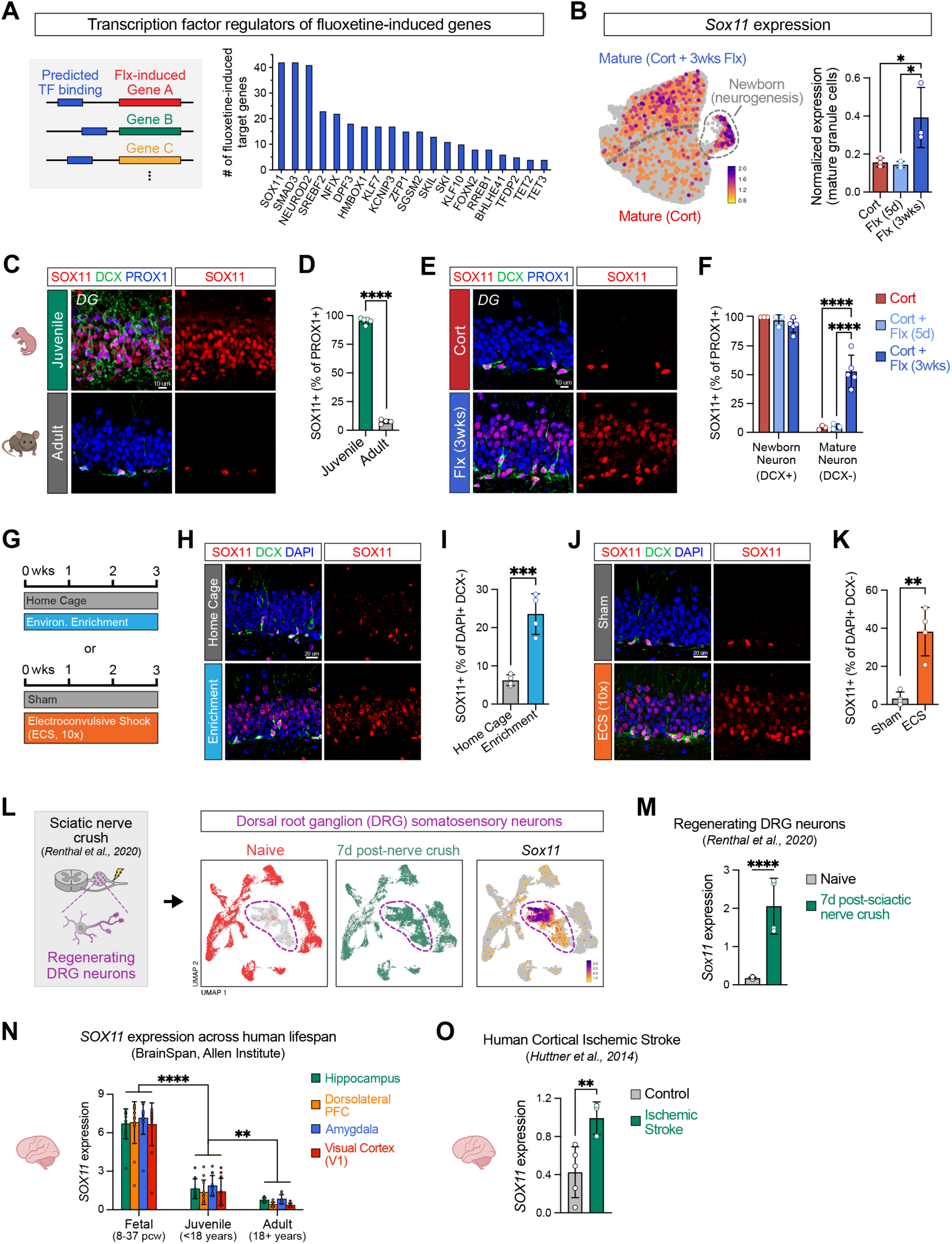
*Sox11* is a conserved developmental switch that is reactivated in mature neurons during physiological contexts and diverse antidepressant interventions. (**A**) Schematic of transcription factors predicted to regulate genes upregulated by 3 weeks of fluoxetine (left). Bar graph of the number of predicted fluoxetine-induced target genes by transcription factor (right; shown are transcription factors upregulated by fluoxetine at p_adj_<0.05). (**B**) Feature plot showing *Sox11* expression (left) and normalized *Sox11* expression selectively in mature granule cells (right; One-way ANOVA, p=0.0282, Tukey’s post-hoc test, n=3 mice/group). (**C-D**) SOX11 protein expression (C) and quantification (D) of SOX11+ granule cells in the DG of juvenile (P10) and adult (∼P120) mice (t-test, p<0.0001, n= 5 mice (juvenile), 4 (adult)). PROX1 marks granule cell neurons. (**E-F**) SOX11 protein expression (E) and quantification (F) of SOX11+ mature (PROX1+ DCX-) and newborn (PROX1+ DCX+) granule cells in the DG of adult mice treated with Cort or Cort + Flx (3wks) (One-way ANOVA, p=0.0002, Tukey’s post-hoc test, n=3 mice (Cort), 3 (Cort + Flx 5d), 5 (Cort + Flx 3wks)). PROX1 marks granule cells, DCX marks newborn granule cells. (**G**) Schematic of environmental enrichment and electroconvulsive shock (ECS) paradigms. Mice received ECS once every other day for a total of 10 sessions (see also *Methods*). (**H-I**) SOX11 protein expression (H) and quantification (I) of SOX11+ mature granule cells in the DG of mice exposed to environmental enrichment or home cage (t-test, p=0.0008, n=4 mice/group). (**J-K**) SOX11 protein expression (J) and quantification (K) of SOX11+ mature granule cells in the DG of mice exposed to ECS or sham (t-test, p=0.0018, n=4 mice/group). (**L-M**) Schematic for transcriptional profiling (*Renthal et al., 2020*)^35^ and *Sox11* expression in regenerating dorsal root ganglion (DRG) neurons (t-test, p<0.0001, n=7 (naive), 4 (7d post-nerve crush). (**N**) *SOX11* expression from bulk RNA sequencing across the human lifespan (BrainSpan, Allen Brain Institute)^75,76^ (Two-way ANOVA, Age p<0.0001, Brain Region p=0.3515, Interaction p=0.9958, Tukey’s post-hoc test, n=15 (fetal), 11 (juvenile), 7 (adult)). (**O**) *SOX11* expression from bulk RNA sequencing in human cortical stroke samples (*Huttner et al., 2014*)^77^ (t-test, p=0.0039, n=5 samples/group). Data = mean ± SD. \**p < 0.05*; \*\**p < 0.01*; \*\*\*\**p < 0.0001*

During neurodevelopment, *SOX11* is among the most highly enriched neuronal genes in both humans and rodents^40,41^ (**Fig. S3B**). In the adult CNS, *Sox11* expression is typically suppressed, but its overexpression is sufficient to induce an immature and highly plastic neuronal state in DG granule cells^42,43^ and promote axonal regeneration in damaged neurons^36,38^. Consistent with this, we found that SOX11 was expressed in nearly all GCs in the neonatal DG but was sharply reduced in the adult DG (**Fig. 3C-D**). Strikingly, after 3 weeks of Flx, SOX11 was reactivated in a spatially distinct subset of mature GCs, an effect that was not overserved after 5 days of Flx (**Fig. 3E-F**). These SOX11+ mature GCs were dissociable from adult neurogenesis and could be distinguished from SOX11+ newborn GCs by their distinct localization in the outer granule cell layer and by their lack of doublecortin (DCX) expression, a marker of newborn GCs. Taken together, these results identify *Sox11* as a developmental switch that is reactivated in mature GCs by chronic Flx treatment.

Our findings raised the possibility that *Sox11* reactivation reflects a more general form of plasticity. To assess this, we examined single cell and bulk RNA-seq datasets from diverse physiological contexts (**Fig. S4** and **Table S3**^31,35,42,44–46^). As expected, *Sox11* was among the top genes enriched in neonatal DG GCs^31^ and in newborn GCs generated through adult hippocampal neurogenesis^42^. *Sox11* also ranked among the top genes in contexts of axonal regeneration and structural plasticity, including in regenerating somatosensory neurons, where it was among the most upregulated genes one week after spinal cord injury^35^, and in DG mossy fiber sprouting^44^, a seizure-induced state of axonal sprouting in GCs. Strikingly, *Sox11* was also enriched in mature GCs under physiological conditions such as environmental enrichment^45^ and downregulated in the hippocampus during aging^46^, suggesting that learning engages a Sox11-mediated neuronal rejuvenation program that is sensitive to the aging process. Notably, we also observed *Sox11* reactivation across multiple antidepressant-like interventions such as ketamine^47^ and electroconvulsive shock^48^ (**Fig. S4B** and **Table S3**^47,48^).

To examine these findings *in situ*, we performed immunostaining for SOX11 in the DG of mice exposed to environmental enrichment or repeated electroconvulsive shock (**Fig. 3G**). In both situations, SOX11 was significantly increased in a spatially distinct subset of DCX-mature GCs that were separate from the subgranular zone where newborn GCs are located (**Fig 3H-K**). Taken together, these results suggest that diverse antidepressant-like modalities could converge on the reactivation of a *Sox11*-mediated developmental plasticity program.

To determine if these interventions also reactivated a developmental-like transcriptional state, we generated a second snRNA-seq dataset following repeated electroconvulsive shock, an antidepressant-like paradigm that rescued behavioral deficits induced by chronic Cort treatment (**Fig. S5**), and analyzed a previously published snRNA-seq dataset of the DG after environmental enrichment^45^ (**Fig. S6**). Both interventions increased the “Developmental Plasticity Score” specifically in mature GCs (**Fig. S5K** and **S6B**), indicating that distinct antidepressant-like modalities converge on the reactivation of developmental-like plasticity programs, separate from their known effects on adult neurogenesis.

We next asked whether *Sox11* reactivation represents a broader developmental switch beyond the hippocampus and whether this mechanism is conserved in the human brain. Neurons in the PNS can re-enter a developmental-like state following nerve injury to facilitate axon regeneration^33–35^. To test whether *Sox11* is re-engaged in this context, we analyzed a previously published snRNA-seq dataset of regenerating dorsal root ganglion neurons after nerve injury^35^ and found that *Sox11* was robustly upregulated (**Fig. 3L-M** and **Fig. S4A**), consistent with its established role as a “regeneration associated gene”^36–38^. In the human brain, we found that *SOX11* expression was prominent across multiple brain regions during neurodevelopment, including areas with well-characterized critical periods such as visual cortex, and was progressively suppressed across the lifespan (**Fig. 3N**), mirroring our findings in the mouse DG.

Ischemic stroke represents another context in which latent plasticity programs are reactivated to support functional recovery after injury^49^. Strikingly, analysis of a transcriptomic dataset from human cortical ischemic stroke samples revealed a significant upregulation of *SOX11* (**Fig. 3O**). Consistent with this, clinical studies have found that Flx and electroconvulsive therapy can enhance motor recovery after ischemic stroke^50^ and in Parkinson’s disease^51^, raising the possibility that analogous *SOX11*-mediated plasticity mechanisms are engaged in humans during antidepressant treatment. Altogether, these findings suggest that the re-engagement of a *Sox11*-mediated developmental program reflects a fundamental form of plasticity that can be recruited not only by pharmacological manipulations but also by physiological contexts in both rodents and humans.

### Sox11 reactivation in mature neurons is necessary for the antidepressant-like behavioral effects of chronic fluoxetine treatment

We next asked whether SOX11 reactivation and rejuvenation in mature GCs was required for the antidepressant-like behavioral effects of chronic Flx treatment. The DG plays a crucial role in affective processing^52–54^ and in mediating the antidepressant-like effects of SSRIs in rodents^5,22,28,29,55^, yet the distinct functional contributions of mature versus newborn GCs have been difficult to dissect, in part because tools to selectively target these populations are limited. To delineate the behavioral relevance of SOX11 reactivation, we generated a mouse model with conditional *Sox11* deletion specifically in mature GCs, while sparing newborn GCs, by crossing *Sox11^flox/flox^*mice with a novel mature GC-specific CreERT2 knock-in line (Dock10-creER; **Fig. 4A-C** and **Fig. S7**).

**Fig 4.**
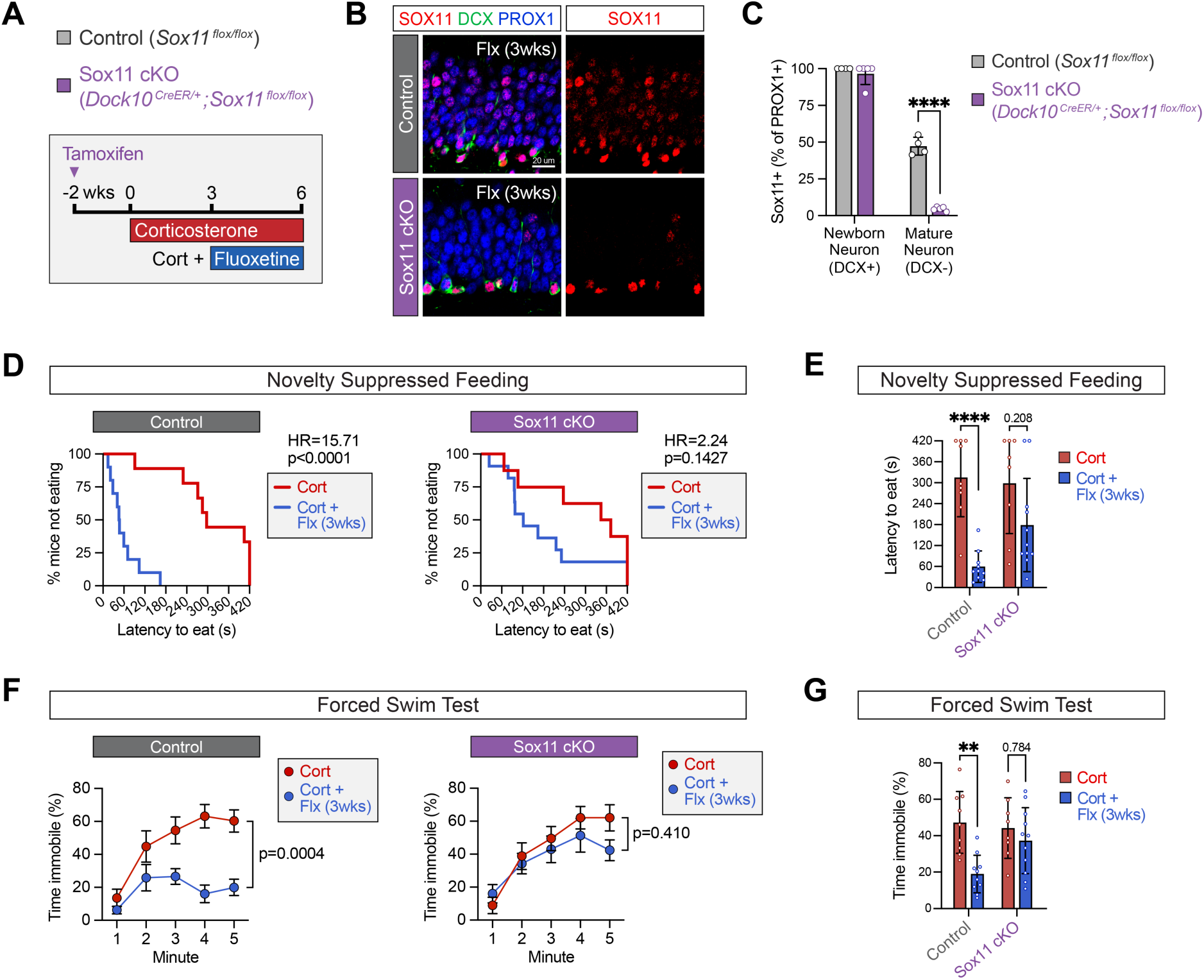
Sox11 reactivation in mature neurons is necessary for the antidepressant-like behavioral effects of chronic fluoxetine treatment. (**A**) Pharmacological and genetic approach. All groups were treated with tamoxifen. (**B**) SOX11 protein expression in the DG of Sox11 cKO mice after 3 weeks of fluoxetine. (**C**) Quantification of SOX11+ mature (PROX1+ DCX-) and newborn (PROX1+ DCX+) granule cells per mouse (t-test, p<0.0001, n=4 mice (Control), 5 (Sox11 cKO). (**D**) Latency to feed in NSF per mouse shown as survival curve in Control (Log-rank Mantel-Cox test, Hazard Ratio=15.71, χ2=17.23, p<0.0001, n=9 (Control-Cort), 10 (Control-Flx)) and Sox11 cKO (Log-rank Mantel-Cox test, Hazard Ratio=2.24, χ2=2.46, p=0.1427, n=8 (Sox11 cKO-Cort), 11 (Sox11 cKO-Flx)). (**E**) NSF data as bar plot (Two-way Cox proportional hazards model, Treatment p<0.0001, Genotype p=0.9724, Interaction p=0.0428, Holm-Sidak post-hoc test, n= same as (D)). (**F**) Immobility time in FST by minute in Control (Two-way RM ANOVA, Treatment p=0.0004, Minute p<0.0001, Interaction p=0.0123, Sidak post-hoc test, n=9 (Control-Cort), 10 (Control-Flx)) and Sox11 cKO (Two-way RM ANOVA, Treatment p=0.4100, Minute p<0.0001, Interaction p=0.1941, n=8 (Sox11 cKO-Cort), 11 (Sox11 cKO-Flx)). (**G**) FST data as bar plot (Two-way ANOVA, Treatment p=0.0017, Genotype p=0.1448, Interaction p=0.0459, Tukey’s post-hoc test, n= same as (F)). Data = mean ± SD (bar plots) and SEM (line plots). \**p < 0.05*; \*\**p < 0.01*; \*\*\**p < 0.001*; \*\*\*\**p < 0.0001*

In Sox11 cKO mice (*Sox11^flox/flox^*; *Dock10^creER/+^*) with mature GC-specific deletion of *Sox11*, we found that the effects of Flx were significantly attenuated in the novelty suppressed feeding and forced swim tests (**Fig. 4D-G**), indicating that *Sox11* reactivation in mature GCs is necessary for these antidepressant-like behavioral effects of chronic Flx treatment. Importantly, Sox11 cKO mice showed normal levels of newborn GCs in response to Flx treatment (**Fig. S7F-G**), indicating that these behavioral effects were not caused by a change in neurogenesis. In the elevated plus maze (EPM), we did not observe differences in open arm exploration or in general locomotion (**Fig. S7H-I**). Collectively, these findings reveal a Sox11-mediated rejuvenation program in mature GCs that is required for several antidepressant-like behavioral effects of chronic Flx treatment.

### Sox11 drives critical period-like structural plasticity in the adult DG following chronic fluoxetine treatment

We next asked whether the cellular mechanisms by which *Sox11* mediates the antidepressant-like effects of Flx are related to Flx-induced critical period-like plasticity. Leveraging our transcription factor analysis (**Fig. 3A**), we examined predicted SOX11 target genes among those upregulated by chronic Flx in mature GCs (**Fig. 5A**). Many SOX11 targets overlapped with genes normally enriched in neonatal relative to adult GCs^31^, consistent with reprogramming to a developmental-like state. Among these was *Bdnf*, a neurotrophin directly regulated by SOX11^56^ that is essential for the synaptic plasticity underlying critical periods^7,15,57,58^ and antidepressant-like effects of Flx^29,55,59^. Consistent with this, *Bdnf* expression was abundant in neonatal GCs but sharply declined in adulthood (**Fig. 5B-C**), and chronic Flx treatment restored *Bdnf* expression in mature GCs of Cort-treated mice (**Fig. 5D-E**). Altogether, these data suggest that *Sox11* re-engages a developmental-like rejuvenation program in part by reactivating *Bdnf*.

**Fig. 5.**
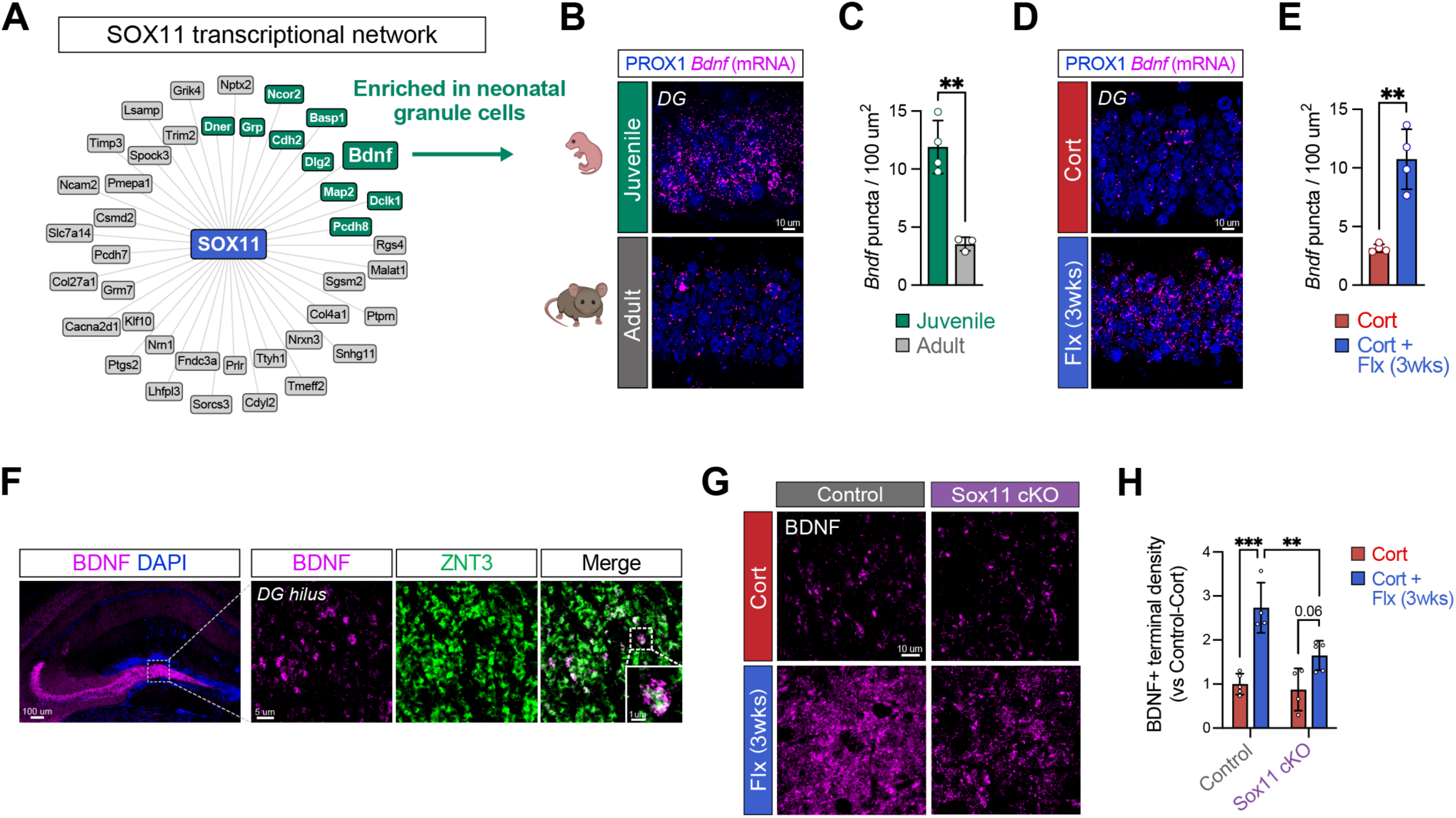
Sox11 drives critical period-like synaptic remodeling in the adult DG following chronic fluoxetine treatment. (**A**) Fluoxetine-induced genes in DG granule cells predicted to be regulated by SOX11. Genes highlighted in green are also upregulated in P5 granule cells relative to P132 at p_adj_<0.05 (Fig. S3A-B). (**B-C**) Image (B) and quantification (C) of *Bdnf* transcript co-labeled with PROX1 in the DG of juvenile (P10) and adult (∼P120) mice (t-test, p=0.0018, n=4 mice (juvenile), 3 (adult)). (**D-E**) Image (D) and quantification (E) of *Bdnf* transcript co-labeled with PROX1 in the DG of mice treated with Cort or Cort + Flx (3 wks) (t-test, p=0.0011, n=4 mice/group). (**F**) BDNF protein expression in the hippocampus. Insets show higher magnification images of BDNF co-labeled with ZNT+ mossy fiber terminals in the hilus as well as a single mossy fiber bouton. (**G**) BDNF protein expression in the DG hilar mossy fiber pathway of Control and Sox11 cKO mice following treatment with Cort or Cort + Flx (3wks). (**H**) Quantification of rejuvenated BDNF+ terminals in the hilar mossy fiber pathway (Two-way ANOVA, Genotype p= 0.0077, Treatment p<0.0001, Interaction p=0.268, Tukey’s post-hoc test, n=4-5 mice/group). Data = mean ± SD. \*\**p < 0.01*; \*\*\**p < 0.001*

BDNF promotes critical period-like synaptic remodeling^7,57,58^ and antidepressant-like effects^29,55,59^, but the specific hippocampal circuits it regulates remain unclear. To investigate this, we immunostained for BDNF and found that within the hippocampus, BDNF protein was highly enriched in the mossy fiber pathway (the axonal projections of GCs) where it co-localized with ZNT3, a marker of mossy fiber terminals^60^ (**Fig. 5F**). To determine if reactivating this critical period-like plasticity was linked to structural plasticity in adulthood, we sparsely labeled mossy fiber terminals and identified the two major terminal types: large mossy terminals and *en* passant varicosities^61^ (**Fig. S8A**). Chronic Flx treatment increased both the number of mossy fiber terminals and the size of these two bouton subtypes (**Fig. S8B-C**), reminiscent of our finding that Flx reactivates an axon regeneration-associated transcriptional program in mature GCs (**Fig. 2G-H**).

To determine whether *Sox11* reactivation in mature GCs was required for this synaptic rejuvenation, we treated mature GC-specific Sox11 cKO and Control mice with chronic Flx and found that Flx-induced increases in rejuvenated synapses (BDNF+) were reduced in Sox11 cKO mice compared to controls (**Fig. 5G-H**). Interestingly, this is consistent with established roles of both SOX11 and BDNF in axon regeneration and structural plasticity after peripheral nerve injury^36–38,62,63^. Altogether, our findings suggest that Flx-induced reactivation of critical period-like plasticity contributes to its antidepressant effects.

### Fluoxetine reinstates a critical period-like environment in the adult DG by remodeling novel extracellular matrix structures

The weeks long latency of Flx’s antidepressant-like effects (**Fig. 1A-C**) and of SOX11-mediated synaptic rejuvenation (**Fig. 3E-F** and **5G-I**) suggest the existence of upstream mechanisms that gate this reactivation. One such candidate is the extracellular matrix (ECM), a poorly understood glycoprotein network that is sparse during early development but progressively accumulates with age, where it acts as a molecular brake to close developmental critical periods^10,64^. In the adult brain, ECM removal and antidepressant treatments can both reopen critical period-like plasticity^11–14,6,7,15,8,16,9^. However, the molecular programs linking ECM remodeling to antidepressant effects are poorly understood, in part because most work has focused on perineuronal nets (PNNs), a specialized subset that constitutes only a small fraction of the total ECM, and on transcriptomic rather than protein-level dynamics.

Consistent with a role in ECM remodeling^17–20^, our transcriptomic results indicated that Flx modifies the ECM composition in the DG (**Fig. 2C**), including upregulation of genes encoding ECM proteinases such as ADAMTS6 and downregulation of structural ECM components such as TENASCIN-R (*Tnr*). To investigate whether ECM structures, including both PNN and non-PNN forms, gate developmental plasticity in the hippocampus, we next examined the ECM landscape *in situ*. Using *wisteria floribunda agglutinin* (WFA), a lectin that binds ECM glycoproteins, we identified canonical PNNs as well as an abundance of non-PNN structures (**Fig. 6A**). While PNNs have been extensively characterized, these other ECM architectures, which comprise the majority of the hippocampal ECM, remain unexplored.

**Fig. 6.**
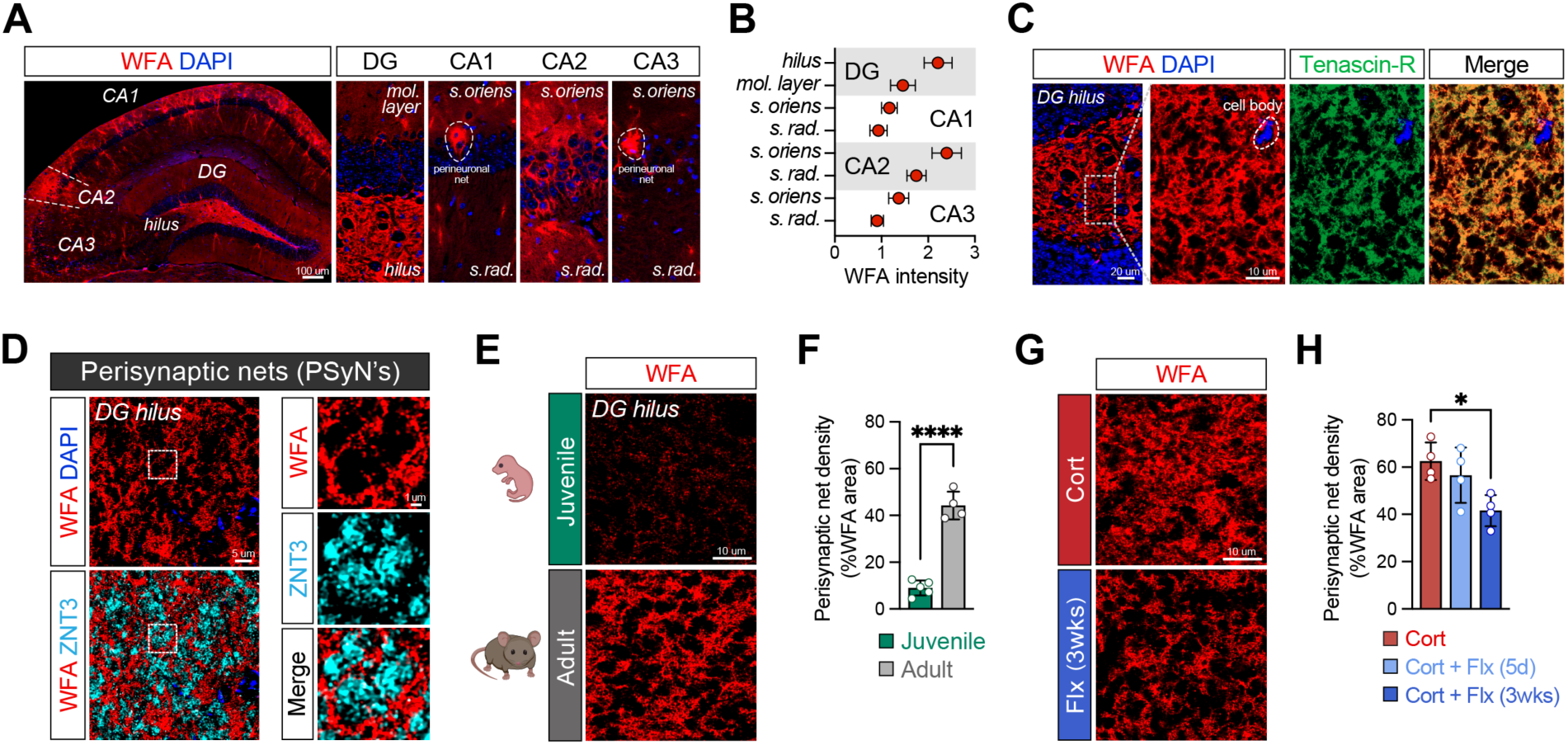
Fluoxetine reinstates a developmental-like environment in the adult DG by remodeling novel extracellular matrix structures. (**A**) Representative image of WFA labeling in the hippocampus. Insets show individual subregions with canonical perineuronal nets (PNNs) surrounding neuronal soma (highlighted), where the majority of the ECM is non-PNN. (**B**) Quantification of WFA intensity in hippocampal subregions. (**C**) ECM in the DG hilus forms porous net-like structures (“perisynaptic nets”) that are immunopositive for WFA and Tenascin-R (TNR). (**D**) Holes in perisynaptic nets (PSyN’s) encase ZNT3+ mossy fiber boutons; inset with higher magnification (right). (**E-F**) WFA staining (E) and quantification (F) of WFA density in DG hilus of juvenile (P10) and adult (∼P120) mice (t-test, p<0.0001, n= 5 mice (juvenile), 4 (adult)). (**G-H**) WFA staining (G) and quantification (H) of WFA density in DG hilus of adult mice treated with Cort or Cort + Flx (3wks) (One-way ANOVA, p=0.0247, Tukey’s post-hoc test, n=4 mice/group). Data = mean ± SD. \**p < 0.05*

WFA labeling was most enriched in CA2 and in the hilus of the DG (**Fig. 6B**), where rejuvenated mossy fiber terminals were densely concentrated following chronic Flx treatment (**Fig. 5F-I**). In the hilus, both WFA and TENASCIN-R labeled a porous net-like structure that encased discrete bundles of ZNT3+ mossy fiber terminals (**Fig. 6C-D**). We termed these expansive structures “perisynaptic nets” (PSyNs) to distinguish them from canonical PNNs that are typically found encasing the soma of interneurons (**Fig. 6D**).

PSyNs were perforated by discrete bundles of synaptic terminals, suggesting a potential role in constraining or sculpting synaptic remodeling. To investigate if PSyNs function as a developmental switch, we quantified their abundance across age. PSyNs were sparse in neonatal mice and expanded significantly in adults, consistent with a role in constraining developmental plasticity programs (**Fig. 6E-F**). Three weeks of Flx treatment reduced PSyN density in the DG of adult mice, suggesting that Flx reinstates a developmental-like environment in the adult brain that is more permissive for structural remodeling (**Fig. 6G-H**). No changes were observed after 5 days of Flx, mirroring the delayed onset of SOX11-mediated synaptic rejuvenation and aligning with the slow kinetics of ECM turnover^21,65^. Altogether, these findings reveal that chronic Flx remodels previously unrecognized ECM structures and reinstates a developmental-like environment in the adult DG.

### The extracellular matrix is a gatekeeper of Sox11-mediated synaptic rejuvenation and antidepressant-like effects

We next asked if directly remodeling the ECM was sufficient to re-engage synaptic rejuvenation in the adult DG. Intracranial delivery of the enzyme Chondroitinase ABC (ChABC) is commonly used to disrupt ECM proteoglycans, but its effects are transient and lack spatial and cell type specificity. To overcome these limitations, we sparsely injected an AAV encoding Cre-dependent ChABC^66^ into the DG of *Dock10-Cre* mice, enabling sustained removal of PSyNs around mature GCs (**Fig. 7A-B**). Notably, this approach produced robust degradation of PSyNs while leaving PNNs intact (**Fig. 7B**).

**Fig. 7.**
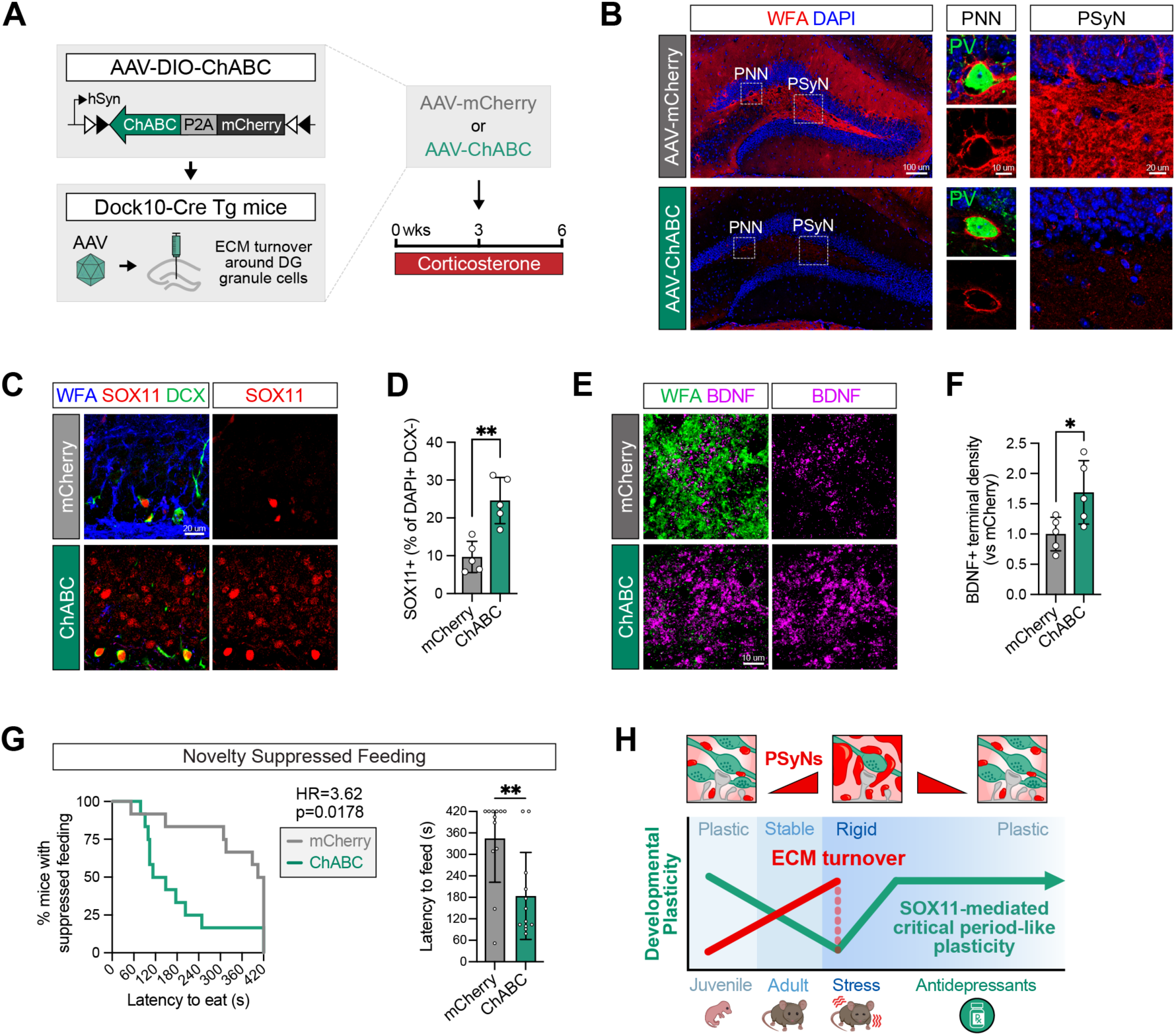
The extracellular matrix is a gatekeeper of Sox11-mediated synaptic rejuvenation. (**A**) Schematic of AAV-based strategy for Cre-dependent expression of Chondroitinase ABC (ChABC) in dentate granule cells and paradigm for Cort treatment. (**B**) Representative images of perineuronal nets (PNNs) surrounding parvalbumin (PV)+ interneurons and perisynaptic nets (PSyNs) in the DG of mice injected with AAV-ChABC or AAV-mCherry. (**C-D**) SOX11 protein expression (C) and quantification (D) of SOX11+ mature granule cells (DCX-) in the DG of mice injected with AAV-mCherry or AAV-ChABC (t-test, p=0.0019, n=5 mice/group). (**E-F**) BDNF protein expression (E) and quantification (F) of BDNF protein levels in the hilar mossy fiber pathway in mice injected with AAV-mCherry or AAV-ChABC (t-test, p=0.0313, n=5 mice/group). (**G**) Latency to feed in NSF per mouse shown as survival curve (left; Log-rank Mantel-Cox test, Hazard Ratio=3.62, χ2=5.62, p=0.0178, n=12 mice/group) and bar plot (right; Mann-Whitney test, p=0.0039). (**H**) Schematic of current model: Perisynaptic nets accumulate with age and chronic stress, limiting plasticity. SSRIs, and possibly environmental enrichment and electroconvulsive shock, remove the ECM to unlock ECM-gated critical period-like plasticity and antidepressant-like effects. Data = mean ± SD. \**p < 0.05*; \*\**p < 0.01*

To investigate whether remodeling the ECM can rejuvenate the DG of chronically stressed mice, we injected AAV-ChABC or a control vector in the DG of Cort-treated animals (**Fig. 7A**). In Cort-treated mice, targeted ECM degradation was sufficient to induce SOX11 expression in DCX-mature GCs (**Fig. 7C-D**), indicating the reactivation of a developmental transcriptional program. This SOX11 reactivation was coupled to an increase in BDNF+ mossy fiber synaptogenesis in the DG (**Fig. E-F**). There was no effect of this manipulation on the levels of newborn GCs (**Fig. S9A-B**), indicating that these effects are not caused by a change in adult neurogenesis.

We next asked whether direct ECM remodeling in the DG was sufficient to induce antidepressant-like effects. In Cort-treated mice, AAV-ChABC and control vector groups showed no significant differences in open arm exploration or overall locomotion in the EPM (**Fig. S9C-D**). By contrast, in the NSF test, AAV-ChABC-injected mice exhibited a reduced latency to feed, similar to the observed effects after chronic Flx treatment (**Fig 7G** and **Fig. S9E**). Altogether, these findings demonstrate that the ECM functions as a modular gatekeeper of synaptic rejuvenation, and that locally remodeling the ECM in the DG is sufficient to recapitulate some of the behavioral effects of chronic Flx treatment (**Fig. 7H**).

## Discussion

Antidepressants, including SSRIs and electroconvulsive therapy, are widely used to treat mood and anxiety disorders, yet their mechanisms of action remain poorly understood. In animal models, both SSRIs and ECM degradation can reopen features of critical period plasticity in the adult brain^11–14,6–8,16,9^, but the cellular substrates and molecular pathways that drive this permissive state and their relevance to antidepressant responses have been unclear. Here, we identify mature DG granule cells as a critical cellular substrate for SSRI-induced plasticity. We find that chronic Flx treatment reactivates SOX11-mediated synaptic rejuvenation in mature GCs to drive antidepressant-like behavioral effects. We further find that this critical period-like synaptic rejuvenation is gated by the ECM, including newly identified perisynaptic nets, and that Flx remodels these ECM structures in the DG to re-engage SOX11-BDNF-mediated structural plasticity and antidepressant-like effects.

Our findings show that Flx induces a strikingly selective transcriptomic response in mature granule cells in the DG. The DG is a uniquely plastic region, in part because it supports the continuous production of newborn GCs through adult hippocampal neurogenesis. While newborn GCs contribute to some but not all behavioral effects of SSRIs^5,22^, our findings reveal that mature GCs also play a distinct and important role in mediating Flx’s antidepressant-like effects. These two populations may act synergistically, consistent with evidence that newborn GCs can modulate the circuit activity of mature GCs^52,67,68^. Importantly, although the extent of adult hippocampal neurogenesis in humans remains debated^69,70^, mature GCs are among the most abundant cell type in both the rodent and human hippocampus^71^, raising the possibility that SSRI efficacy in humans is driven by rejuvenation of mature neurons, independent of adult neurogenesis.

Within the mature GC population, we find reactivation of the developmental transcription factor SOX11 as a critical driver of BDNF-associated structural plasticity and antidepressant-like effects. BDNF has well-established roles in regulating critical period plasticity in areas such as the visual cortex^7,15,57,58^, thus positioning SOX11 as a developmental switch that can be reactivated in mature neurons to reinstate critical period-like mechanisms. This reactivation of developmental plasticity is reminiscent of axon regeneration programs in the PNS, which recruit “regeneration-associated genes” that support axonogenesis, including both *Sox11* and *Bdnf* ^35–39,62,63^. Interestingly, *Sox11* reactivation in mature GCs is not restricted to SSRI treatment, as we also observe it after repeated electroconvulsive shock and environmental enrichment. Taken together, these findings suggest that SOX11-mediated reactivation of developmental plasticity is a fundamental form of plasticity that is harnessed by antidepressants to drive neural circuit remodeling. Consistent with this, clinical studies have found that Flx and electroconvulsive therapy can enhance motor recovery after ischemic stroke^50^ and in Parkinson’s disease^51^, raising the possibility that analogous plasticity mechanisms are engaged in humans during antidepressant treatment.

Chronic stress and diverse antidepressant modalities bidirectionally remodel the ECM^16–20^, but how this drives antidepressant-like behavioral effects remains unclear. This is in part because most studies have focused on perineuronal nets (PNNs), a specialized subset that represents only a small fraction of the total ECM. In this study, we discover hippocampal ECM structures distinct from canonical PNNs, including expansive net-like structures in the DG that envelop clusters of axonal boutons, which we term perisynaptic nets (PSyNs). We find that directly remodeling PSyNs in the DG while sparing PNNs via targeted ECM degradation is sufficient to reactivate SOX11-BDNF-mediated axonogenesis and promote antidepressant-like effects. Interestingly, this mirrors the well-established role of the ECM as an inhibitory barrier to axon regeneration and structural plasticity following nerve injury^72,73^. Taken together, our findings demonstrate that the distinct ECM architecture in the DG functions as a modular gatekeeper of critical period-like rejuvenation in mature neurons and establishes a molecular mechanism linking ECM remodeling in the DG to antidepressant-like effects.

Our results reveal that PSyNs are dynamic structures that expand as the brain matures yet can be robustly remodeled by chronic Flx treatment. The distinctive ECM architecture in the DG may underly the striking and selective transcriptomic reprogramming we observe in mature GCs following Flx treatment. Additionally, the exceptionally long-lived nature of many ECM proteins and the extended time frame required for remodeling the ECM^21,65^ also provides a possible substrate for the weeks long delay in SOX11 reactivation and the antidepressant-like effects of Flx, which mirror the delayed onset of SSRI efficacy in patients. Although ECM networks are long-lived, they remain dynamic through mechanisms such as ECM proteases and microglia-mediated phagocyotosis^74^. Notably, we found that microglia were among the earliest cell types to respond to Flx treatment in our dataset, highlighting neuron-glia-ECM interactions as an important avenue for future investigation (**Fig. 1I**).

Our findings provide new insights for understanding antidepressant efficacy in humans. The ability of Flx and electroconvulsive shock to rejuvenate mature GCs points to a novel mechanism by which antidepressants may enhance neural plasticity through re-engaging developmental transcriptional programs. Interestingly, this mechanism is independent of adult hippocampal neurogenesis and may be particularly relevant in humans, where the extent and functionality of adult neurogenesis remain debated. Better understanding how neurons access latent developmental programs and their effects on neuronal network functions represents a promising avenue for developing more effective therapeutic strategies for psychiatric and neurological disorders.

## Acknowledgments

We are grateful for S. Schacher for helpful comments on this manuscript. We thank S. Tonegawa for sharing the Dock10-cre mouse strain and S. Dudek for sharing the AAV-ChABC and AAV-mCherry viral reagents. We are grateful for A. Santiago and J. Castello-Saval for assistance with electroconvulsive shock and tissue preparation. We thank the Single Cell Analysis Core of the Columbia University Sulzberger Genome Center under the leadership of Dr. E. Bush and the Columbia Stem Cell Initiative Flow Cytometry core facility at Columbia University Irving Medical Center under the leadership of Dr. M. Kissner.

## Author contributions

Conceptualization: PTN, RH; Methodology: PTN, YX, RH; Investigation: PTN, ST, ES, YS; Visualization: PTN, RH; Funding acquisition: PTN, RH; Project administration: PTN, RH; Supervision: RH; Writing – original draft: PTN, RH; Writing – review & editing: all authors

## Data and materials availability

All data needed to evaluate the conclusions in this paper are present in the paper or the supplementary materials. All mouse strains were obtained or generated at Jackson Laboratories except for Dock10-cre Tg line, which was a gift from S. Tonegawa. Viral reagents used in this study were a gift from S. Dudek. Single nuclei RNA-seq data are available through Gene Expression Omnibus (GEO) accession #GSE289328 (Dataset 1) and #GSE289329 (Dataset 2).

---

**Supplemental Fig. S1.**
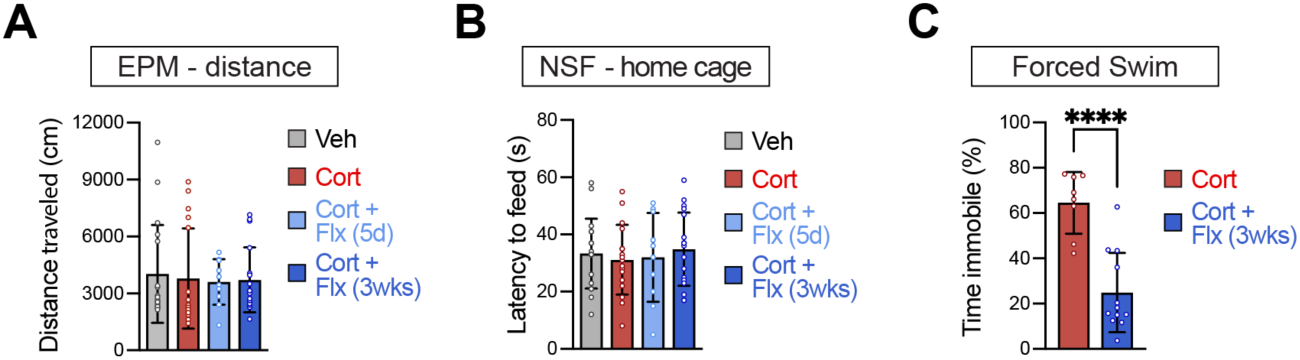
Additional data pertaining to behavioral studies; related to Figure 1. (**A**) Distance traveled in the EPM per mouse (One-way ANOVA, p=0.96, n=18 (Veh), 17 (Cort), 10 (Cort + Flx 5d), 18 (Cort + Flx 3wks)). (**B**) Latency to feed in the home cage (One-way ANOVA, p=0.85, n= same as (A)). (**C**) Immobility time in forced swim test per mouse (unpaired t-test, p<0.0001, n=8 (Cort) and 12 (Cort + Flx 3wks)). Data = mean ± SD.

**Supplemental Fig. S2.**
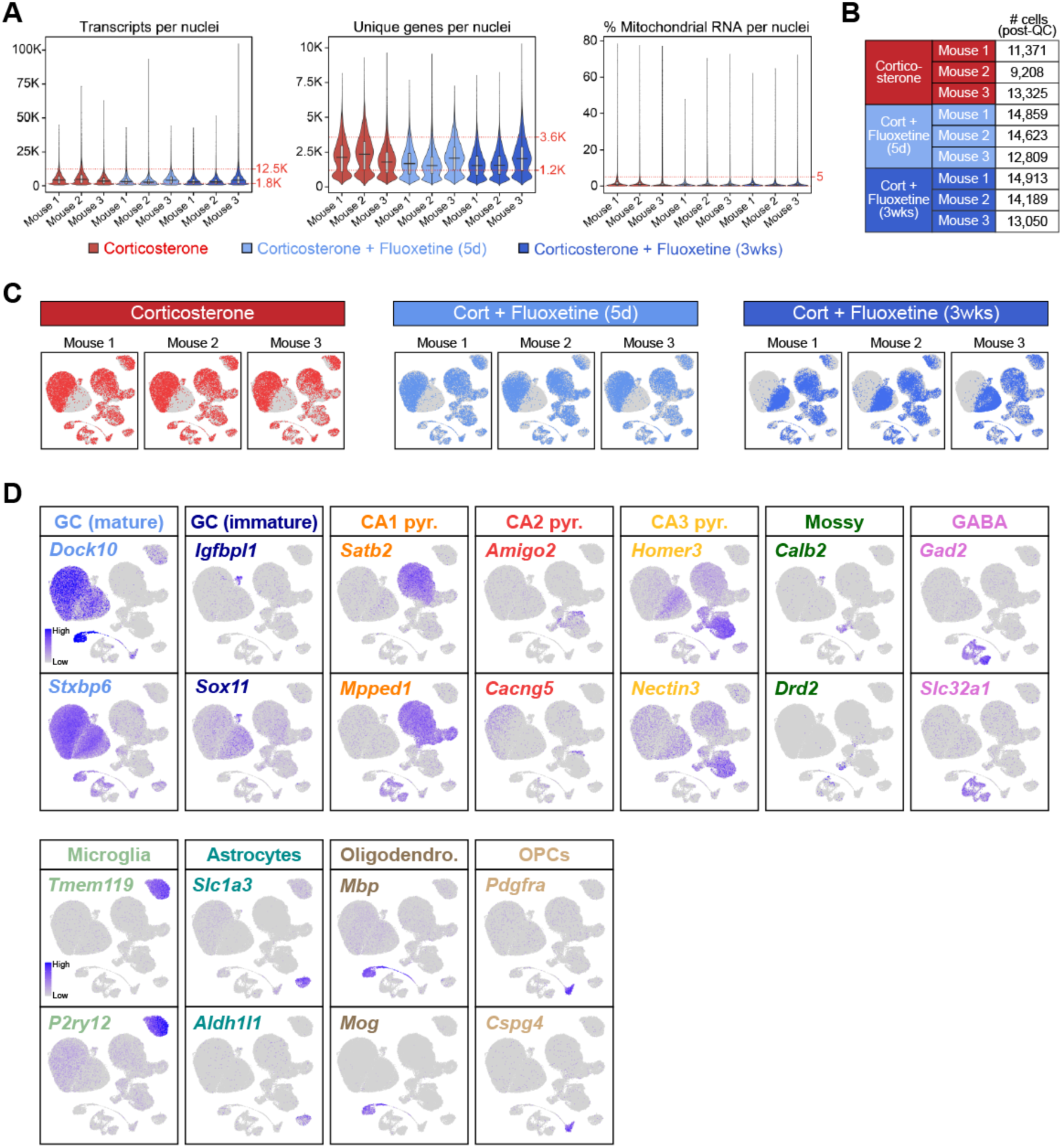
Quality control and validation of snRNA-seq data; related to Figure 1. (**A**) Quality control metrics for single nuclei RNA-sequencing in Cort, Cort + Flx (5d), and Cort + Flx (3wks) groups. Dashed lines indicate minimum and maximum threshold settings. (**B**) Number of nuclei per mouse after quality control thresholding. (**C**) Comparison of biological replicates following unsupervised clustering of hippocampal nuclei. (**D**) Representative markers for major neuronal and glial cell type in the hippocampus. GC = granule cells (dentate gyrus), Pyr. = pyramidal neuron, Oligodendro. = oligodendrocytes, OPCs = oligodendrocyte progenitor cells.

**Supplemental Fig. S3.**
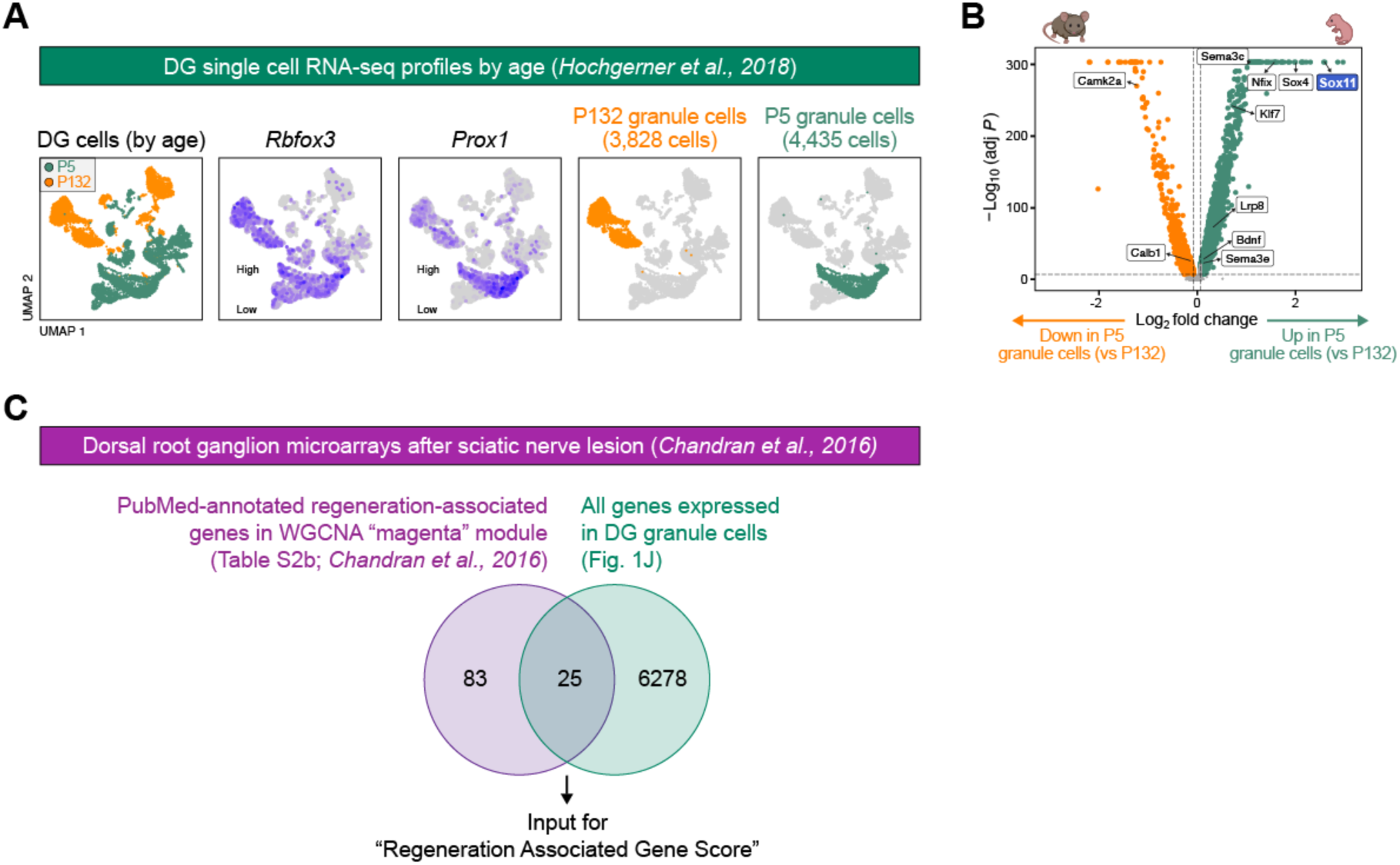
Additional data pertaining to snRNA-seq experiments; related to Figure 2. (**A**) Analysis of single cell RNA-seq dataset^31^ comprised of 8,264 cells from the dentate gyrus from P5 and P132 mice. Unsupervised clustering analysis and identification of granule cells are shown (n=6 pooled mice (P5), 3 male/1 female mice (P132)). (**B**) Volcano plot of differentially expressed genes in P5 granule cells versus P132 granule cells from (A). (**C**) WGCNA-identified genes (“magenta” module^39^) enriched in regenerating dorsal root ganglion neurons following nerve injury from *Chandran et al., 2016*. The overlap between PubMed-annotated genes associated with axon regeneration (Table S2b from *Chandran et al., 2016*) and genes expressed in DG granule cells were used as input for the “Regeneration associated gene” score.

**Supplemental Fig. S4.**
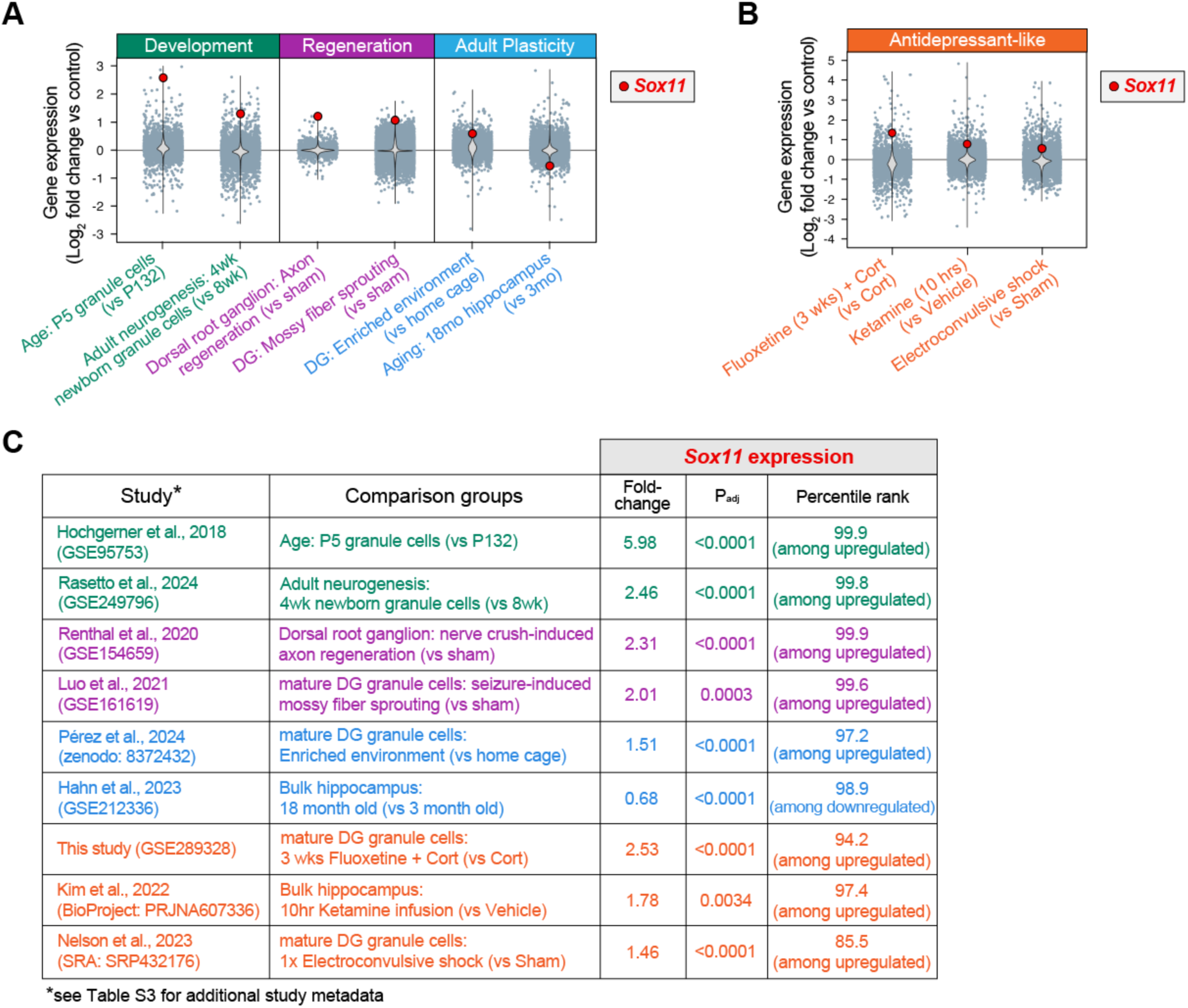
*Sox11* is reactivated in diverse physiological and antidepressant-like modalities. (**A-B**) Log_2_ fold change in gene expression from published single cell or bulk RNA-seq datasets relative to each dataset’s own control. Dots represent individual genes with violin plots illustrating the distribution of expression values; Sox11 is highlighted in red. (**C**) Additional information for each transcriptomic dataset. See **Table S3** for full details of each dataset and experimental conditions.

**Supplemental Fig. S5.**
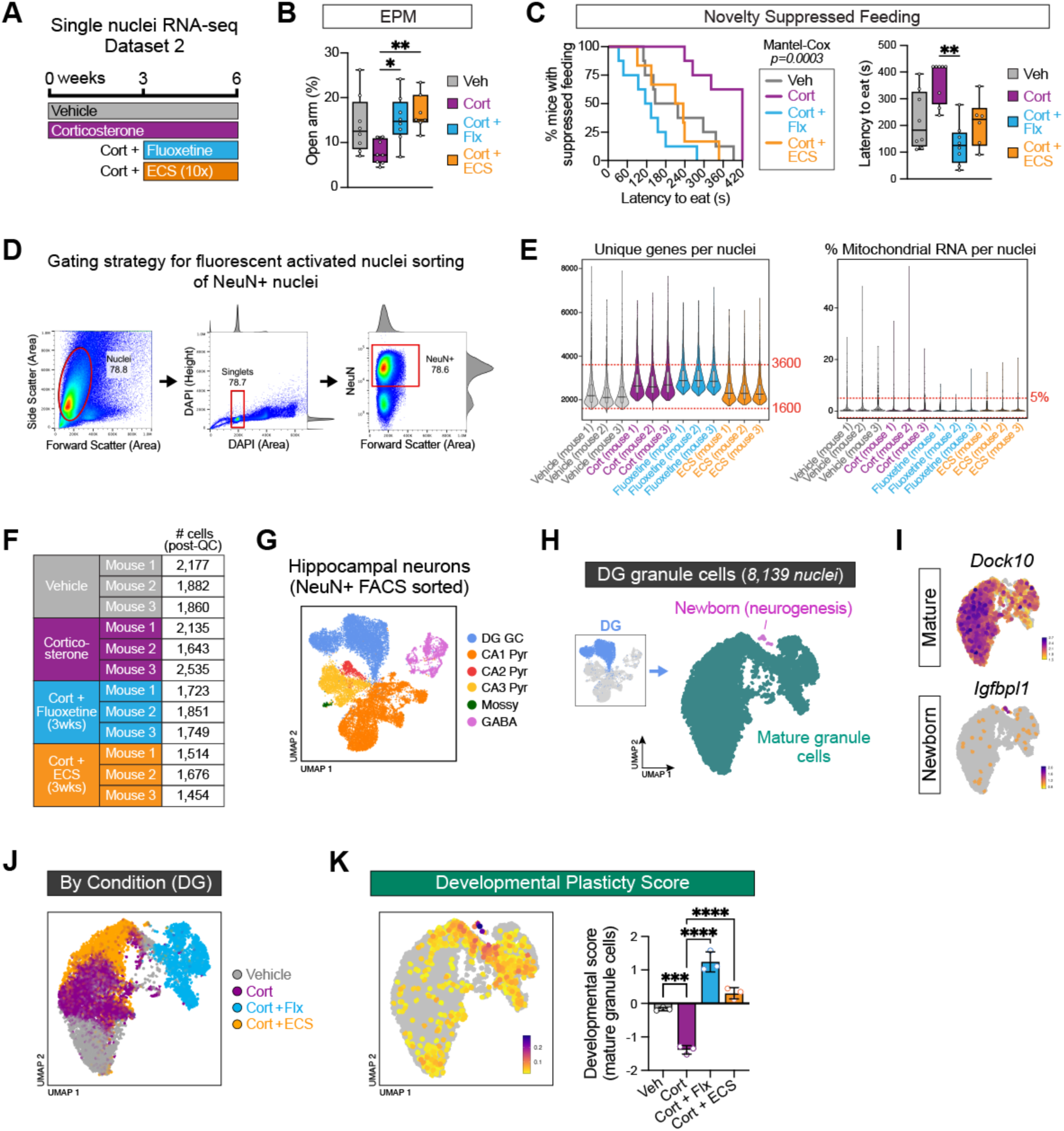
Neurodevelopmental plasticity programs are re-engaged following electroconvulsive shock in an independent snRNA-seq dataset. (**A**) Schematic for pharmacological and electroconvulsive shock (ECS) approach for snRNA-seq of hippocampal nuclei (Dataset 2; see *Methods*). Mice received ECS treatment once every other day for a total of 10 sessions, starting after 3 weeks of Corticosterone treatment. (**B**) Time spent in the open arm of the EPM per mouse (One-way ANOVA, p= 0.0076, Tukey’s post-hoc test, n=8 (Veh), 8 (Cort), 8 (Flx), 7 (ECS)). (**C**) Latency to feed in NSF per mouse shown as survival curve (left; Log-rank Mantel-Cox test, χ2=18.78, p=0.0003) and box plot (right; Kruskal-Wallis ANOVA, p=0.0022, Dunn’s post-hoc test, n=same as (B)). (**D**) Representative gating strategy to isolate NeuN+ neuronal nuclei from the hippocampus using fluorescence activated nuclei sorting for downstream snRNA-seq. (**E**) Quality control metrics for snRNA-seq in Vehicle, Cort, Cort + Flx, and Cort + ECS groups. Dashed lines indicate minimum and maximum threshold settings. (**F**) Table showing number of nuclei per mouse after quality control thresholding. (**G**) Unsupervised clustering (UMAP) of 22,199 single nuclei transcriptomic profiles of hippocampal neurons labeled by cell type. (**H**) Subclustering of mature granule cells from (G). (**I**) Expression of maturation-related marker genes in granule cells. (**J**) Granule cell nuclei labeled by treatment condition: Vehicle (grey), Corticosterone (purple), Cort + Flx (blue), and Cort + ECS (orange). (**K**) Feature plot showing average expression of ‘Developmental Plasticity Score’ (left; derived from Fig. 2D) and z-scores selectively in mature granule cells per mouse (right; One-way ANOVA, p<0.0001, Tukey’s post-hoc test, n=3 mice/group). Barplot Data = mean ± SD. \**p < 0.05*; \*\**p < 0.01*; \*\*\**p < 0.001*; \*\*\*\**p < 0.0001*.

**Supplemental Fig. S6.**
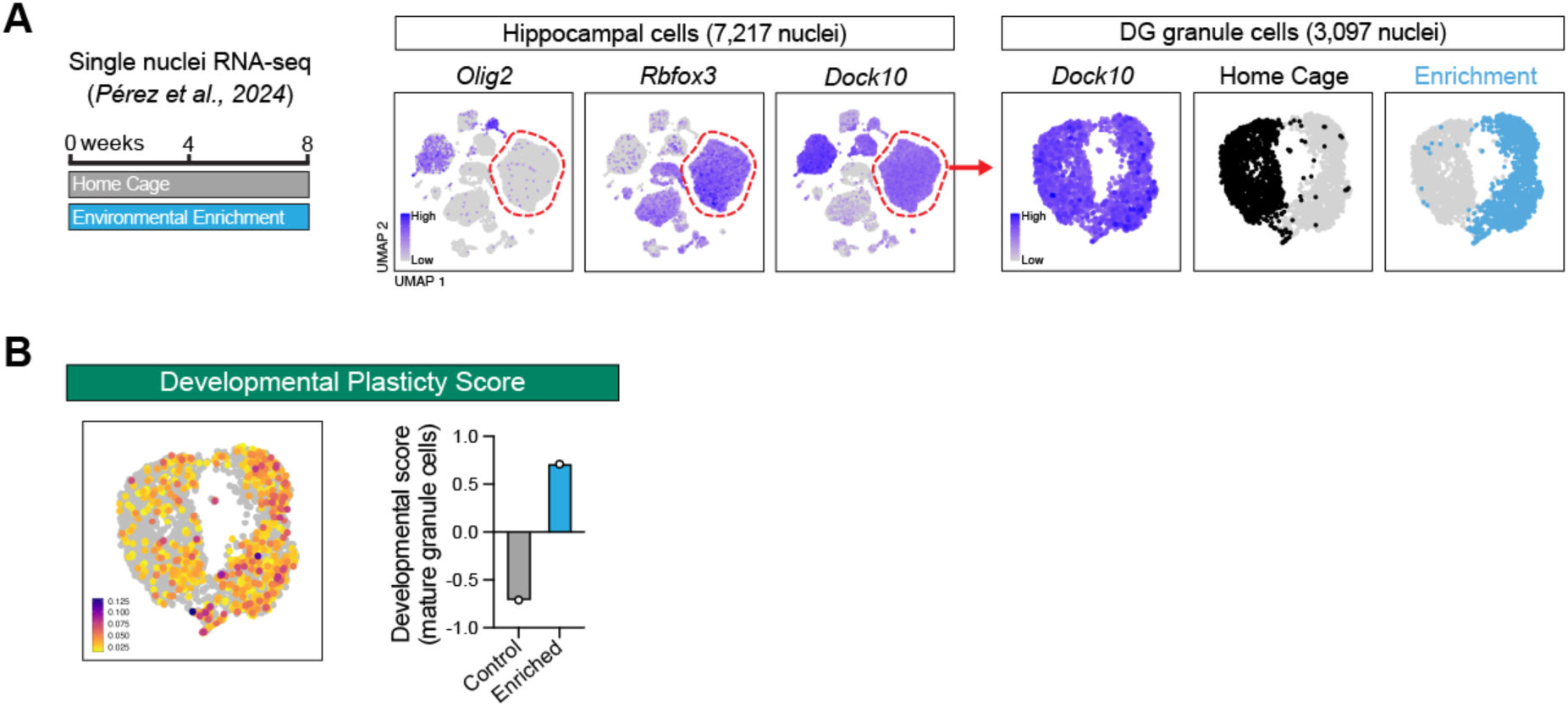
Neurodevelopmental plasticity programs are re-engaged following environmental enrichment. (**A**) Analysis of a snRNA-seq dataset^45^ comprised of 7,217 hippocampal nuclei from 9 week-old mice housed in environmental enrichment (2 months) or control cages. Unsupervised clustering analysis, identification, and subclustering of mature DG granule cells (3,097 nuclei) are shown. (**B**) Feature plot showing average expression of ‘Developmental Plasticity Score’ (left; derived from Fig. 2D) and z-scores in mature granule cells per mouse (right; n=3 mice pooled into 1 replicate per condition).

**Supplemental Fig. S7.**
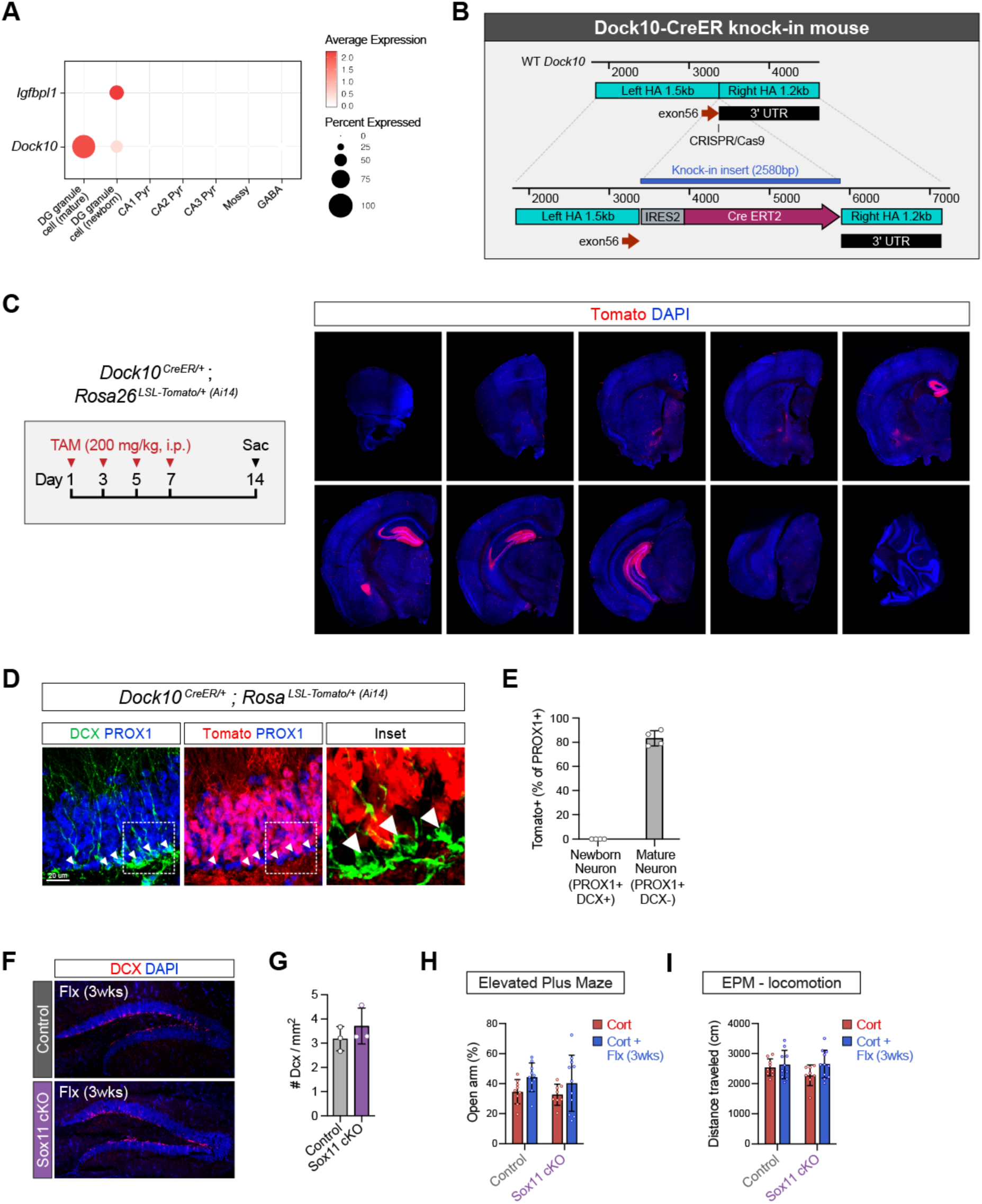
Generation of mature granule cell-specific Dock10^creER^ knock-in mouse line; related to Figure 4. (**A**) Expression of granule cell maturity genes across hippocampal neuronal populations. (**B**) Strategy for the generation of Dock10-creER knock-in mouse line (see also *Methods*). (**C**) Recombination activity across brain regions in adult Dock10-creER mice. (**D**) Specificity of recombination in the DG of Dock10-creER mice. (**E**) Quantification of recombination in mature (PROX1+ DCX-) and newborn (PROX1+ DCX+) granule cells. (**F**) Neurogenesis in Control and Sox11 cKO mice treated with fluoxetine for 3 weeks. (**G**) Quantification of DCX+ newborn neurons (t-test, p=0.3686, n=3 mice/group). (**H**) Time spent in the open arm of the EPM per mouse (Two-way ANOVA, Treatment p=0.0379, Genotype p=0.4604, Interaction p=0.7957, Tukey’s post-hoc test, n=9 (Control-Cort), 10 (Control-Flx), 8 (Sox11 cKO-Cort), 11 (Sox11 cKO-Flx)). (**I**) Distance traveled in the EPM per mouse (Two-way ANOVA, Treatment p=0.0857, Genotype p=0.3640, Interaction p=0.3044, n=9 (Control-Cort), 10 (Control-Flx), 8 (Sox11 cKO-Cort), 11 (Sox11 cKO-Flx)).

**Supplemental Fig. S8.**
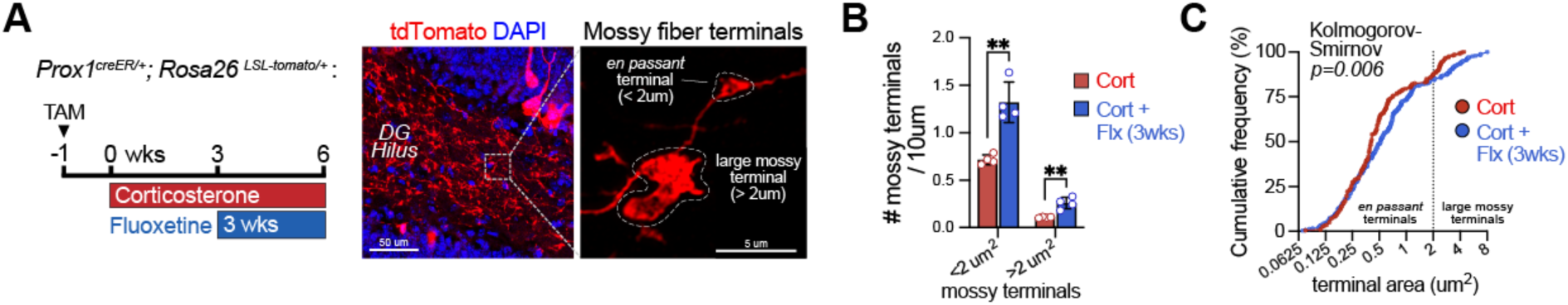
Chronic fluoxetine treatment drives critical period-like axonal remodeling and structural plasticity in the adult DG; related to Figure 5. (**A**) Genetic and pharmacological approach to sparsely label mossy fiber terminals (left) and representative images of large mossy terminals and *en passant* varicosities in the hilus (right). (**B**) Quantification of hilar mossy fiber terminal numbers in mice treated with Cort or Cort + Flx (3wks) (t-test, p=0.0015 (<2 um^2^), p=0.0028 (>2 um^2^), n= 4 mice/group). (**C**) Quantification of mossy fiber terminal area as cumulative distribution in mice treated with Cort or Cort + Flx (3wks) (Kolmogorov-Smirnov test, p=0.0063, n= 200 terminals (Cort), 245 (Cort + Flx 3wks)). Data = mean ± SD.

**Supplemental Fig. S9.**
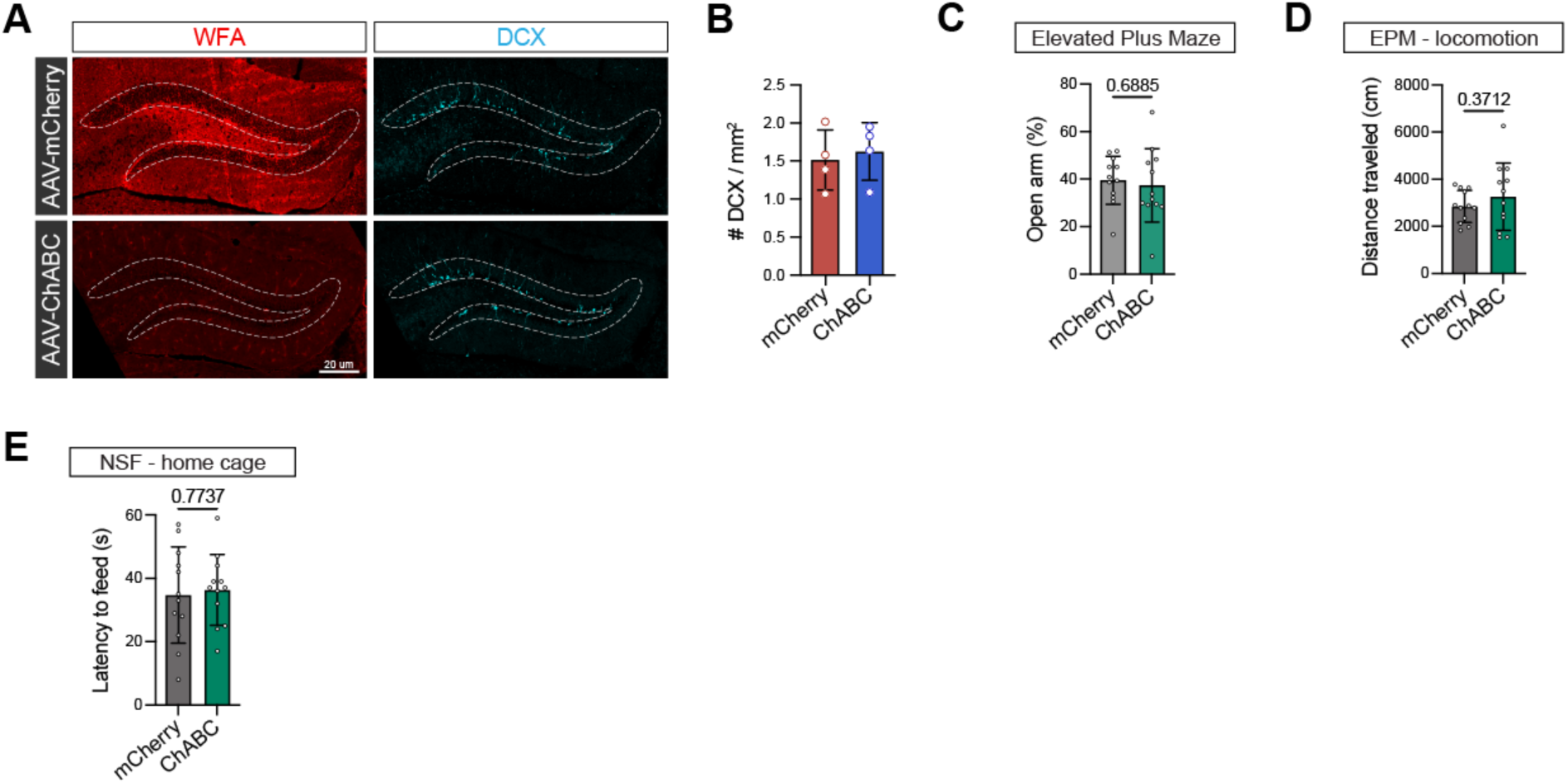
Additional data pertaining to AAV-ChABC studies; related to Figure 7. (**A**) Neurogenesis in mice injected with AAV-mCherry or AAV-ChABC treated with Cort for 3 weeks. (**B**) Quantification of DCX+ newborn neurons (t-test, p=0.6966, n=4 mice/group). (**C**) Time spent in the open arm of the EPM (t-test, p=0.6885, n=12 mice/group). (**D**) Distance traveled in the EPM in mice injected with AAV-mCherry or AAV-ChABC (t-test, p=0.3712, n=12 mice/group). (**E**) Latency to feed in home cage in mice injected with AAV-mCherry or AAV-ChABC (t-test, p=0.7737, n=12 mice/group).

## Materials and Methods

### Animal Studies

All mouse strains were maintained in the New York State Psychiatric Institute specific pathogen-free facility and maintained on a 12-hour light/dark schedule. Mouse protocols were approved by the Institutional Animal Care and Use Committee of Columbia University and the Research Foundation for Mental Hygiene, Inc. and were conducted in accordance with the NIH Guide for the Care and Use of Laboratory Mice. Care was taken to minimize the number of mice used and their suffering.

Littermate controls and both male and female mice were used for all experiments whenever feasible. All mice were maintained on a C57BL/6 background. *Sox11^flox/flox^* (strain #035427), *Prox1-CreER^T^*^2^ (strain #022075) and *Ai14* (strain #007914) mice were purchased through Jackson Labs. *Dock10-cre* transgenic mice were a generous gift from Dr. Susumu Tonegawa^78^.

For environmental enrichment studies, mice were housed in large ventilated rat cages containing standard nesting material, rodent foraging toys, and one running wheel per cage. Nesting and foraging toys were replaced every 3 days throughout a 3 week enrichment period. Control mice were maintained under standard housing conditions.

### Generation of mature granule cell-specific CreER^T2^ knock-in mouse line

*Dock10-IRES-CreER^T^*^2^ knock-in mice were generated on a C57BL/6J background by targeted insertion of an IRES-CreERT2 cassette at the 3′ end of the endogenous *Dock10* locus (transcript ENSMUST00000077946.12). CRISPR/Cas9-mediated genome editing was used to insert the IRES-CreERT2 sequence immediately downstream of the *Dock10* stop codon to enable tamoxifen-inducible Cre recombination under control of the native *Dock10* regulatory elements (see also Fig. S7). Genome editing was performed in C57BL/6J zygotes (JAX strain #000664) at Jackson Labs and founders were bred to C57BL/6J mice to establish germline transmission. Founder and N1 animals were screened for correct targeting by long-range PCR across the insertion junctions followed by sequencing. N1 animals with the correctly targeted knock-in allele were subsequently bred to C57BL/6J mice to establish the line. *Dock10-IRES-CreER^T^*^2^ knock-in mice will be made available through Jackson Labs.

### Pharmacological Approach

Corticosterone (Sigma, C2505) was dissolved at a concentration of 35 µg/mL in autoclaved deionized water containing 2 mg/mL beta-cyclodextrin (Chem-Impex, 32075) to improve solubility. The solution was then incubated in a 37°C water bath sonicator for 30 minutes to ensure complete dissolution. Fluoxetine hydrochloride (TCI Chemicals, F0750) was added at a final concentration of 160 ug/mL under the same conditions. Drug solutions were delivered ad libitum in drinking water in opaque bottles to protect from light. Fluoxetine concentrations were previously optimized to achieve brain pharmacokinetics equivalent to 18 mk/kg/day^79^. Tamoxifen (Sigma, T5648) was dissolved in Corn Oil (Sigma-Aldrich) by incubation in a 37°C water bath with sonication and administered by intraperitoneal injection at 200 mg/kg. Injections were administered four times in total, every other day. Tamoxifen solutions were prepared fresh for each injection.

### Behavioral Assays

Anxiety and depressive-like behaviors were assessed using the elevated plus maze (EPM), novelty suppressed feeding (NSF), and forced swim tests (FST). Male and female mice (2-4 months old) on a C57BL/6 background were used for all behavioral assays. Testing and analyses were performed blinded to genotype and treatment, and conditions were counterbalanced across sex and littermates. All behavioral experiments were initiated in the morning (local Eastern Standard Time, EST).

For both the EPM and NSF tests, mice were tested in 7 minute sessions under low light conditions. Ambient room lights were turned off, and lamps were positioned to provide targeted illumination (∼150 lux) over the open arms (EPM) or the center of the arena (NSF). In the NSF, mice were food deprived for 24 hours prior to testing. Behavior was recorded using Logitech webcams with OBS Studio software. Time spent in the open arms during the EPM was quantified using automated location tracking with the open-source ezTrack pipeline^80^. Latency to feed in the NSF, defined as the amount of time to first bite the food pellet in the center of the area, was scored by live observation and assisted with video recordings. To measure home cage feeding latency, mice were first placed in a holding cage and then returned sequentially to their respective home cages. The latency to bite a food pellet placed at the center of the home cage was measured during a 5 minute session.

For the FST, mice were placed individually into a 3 L beaker containing room temperature water and recorded for 5 minutes. Immobility time was quantified using the automated freeze analysis module in ezTrack.

### Viral Reagents and Stereotaxic Injections

AAV-hSyn-DIO-ChABC-2a-mCherry and AAV-hSyn-DIO-mCherry viral reagents were a generous gift from Dr. Serena Dudek^66^ and packaged by the National Institute of Environmental Health Sciences viral vector core facility at titers of 4e12 and 1e13 GC/ml, respectively.

All stereotaxic injections were performed with a Kopf stereotaxic apparatus (David Kopf, Tujunga, CA) with a microdispensing pump (World Precision Instruments) holding a 33g beveled NanoFil needle (World Precision Instruments, NF33BV). Mice were anesthetized with 1.5% isoflurane at an oxygen flow rate of 1L/min, headfixed with a stereotaxic frame, and treated with ophthalmic eye ointment. Fur was shaved and the incision site was sterilized with 70% ethanol and Betadine prior to surgical procedures. Craniotomy was performed using a surgical drill. To induce turnover of extracellular matrix, 1.5uL of AAV-hSyn-DIO-ChABC-2a-mCherry (diluted 1:100) or AAV-hSyn-DIO-mCherry (diluted 1:1000) viral reagents were bilaterally injected into the dentate gyrus (−3mm AP, ±2.4mm ML, and −2.75mm DV) at a rate of 3 nL/sec. Body temperature was maintained throughout surgery using a heating pad. Post-surgery, Carprofen (Covetrus, 059150) was administered as needed by intraperitoneal injection at a concentration of 5 mg/kg.

### Single Nuclei RNA Sequencing

#### Tissue Isolation and Processing

Whole and intact hippocampi were isolated under a dissecting microscope by separating the cerebral cortex from underlying diencephalon and gently scooping the hippocampus from the overlying cortex with a microspatula. Excess myelin and meninges were removed with forceps. Hippocampi (2 per mouse) were mechanically dissociated using a 2 mL glass tissue homogenizer (Kontes Glassware) on ice using twelve strokes of the tight pestle in 1% BSA-PBS. Nuclei were filtered using a 40 um strainer and pelleted at 300g for 5 minutes at 4°C in a centrifuge. Nuclei were then resuspended in 22% Percoll (GE Healthcare) and centrifuged at 900 g for 20 minutes with acceleration set to 4 and deceleration set to 1 to separate cellular debris. The debris-containing supernatant was removed, and nuclei were washed and resuspended in 1% BSA-PBS before being pelleted at 300 g for 5 minutes at 4°C. Nuclei were resuspended in 1% BSA-PBS prior to loading into the 10x Chromium controller platform. All buffers contained Protector RNase Inhibitor (Roche) at 0.2 U/uL.

Nine samples were prepared corresponding to 3 mice per group: Corticosterone, Corticosterone + Fluoxetine (5d), and Corticosterone + Fluoxetine (3wks). All tissue isolation and library preparation steps were performed on the same day to mitigate batch effects. Three to four months old C57BL/6 male mice were used for all sequencing studies.

#### Library Preparation and Sequencing

Hippocampal nuclei were sequenced using the 10x Single Cell Gene Expression 3’ (v3) platform. Approximately 15,000-20,000 intact nuclei from each biological replicate were loaded into each well of a Chromium Chip A and combined into droplets with barcoded gel beads using the Chromium controller according to the manufacturer instructions. Libraries were prepared by the Single Cell Analysis Core of the Columbia University Sulzberger Genome Center following manufacturer instructions and sequenced on an Illumina Novaseq 6000. Libraries were sequenced to a mean depth of ≥20,000 reads per nucleus with ≥35% sequencing saturation. Reads were processed using the Cell Ranger v6.1.2 pipeline and aligned to the mm10-2020-A mouse reference genome. Approximately 140K total cells were recovered prior to quality control processing.

#### Quality Control and Data Analysis

Following alignment in Cell Ranger, molecular counts data were imported into R (v4.3.0) for quality control, clustering, and differential expression analysis using Seurat^81^ (v4.4.0). Nuclei with <1,200 or >3,600 detected genes per nuclei, or >5% mitochondrial reads, were excluded prior to downstream analyses. Genes detected in fewer than five nuclei were also removed. After quality control filtering, 9,000-15,000 nuclei were retained for each biological replicate (119,346 total nuclei). See also Figure S2.

Counts data underwent normalization and variance stabilization using SCTransform in Seurat, regressing out percent mitochondrial reads and total counts per cell. The top 6,000 most variable genes were used to calculate 50 principal components (PCs), and the top 35 PCs were used for nearest neighbor graph construction, UMAP, and clustering. UMAP plots for all hippocampal cell types were constructed using RunUMAP with n.neighbors = 45 and min.dist = 0.7. Cell types were assigned using expression of canonical cell type-specific marker genes. Rare populations (e.g., epithelial cells and pericytes) were identified but excluded from downstream analyses due to low cell numbers. Data visualization was performed using Seurat and the scCustomize package^83^.

DG granule cells were isolated based on expression of the granule cell markers *Prox1*, *Dock10*, and *Stxbp6*^82,84^, yielding 34,976 nuclei. Because oligodendrocyte lineage cells share several granule cell marker genes (e.g. *Prox1*, *Dock10*, and *Stxbp6*), these cells were excluded by positive identification of oligodendrocyte markers (*Olig2*, *Mbp*, *Mog*) and oligodendrocyte progenitor cell markers (*Pdgfra*, *Cspg4*). The top 6,000 most variable genes were then used to recalculate PCs, generate UMAP embeddings, and define clusters. UMAP plots for DG granule cells were using RunUMAP with n.neighbors = 35 and min.dist = 0.8. Clusters were determined using a resolution of 0.1 using the ClusTree package^85^. For UMAP visualizations, granule cell nuclei were integrated across biological replicates using Harmony^86^ on the top 20 PCs.

Differential gene expression analyses were conducted using FindMarkers with the MAST test in Seurat. For analyses of DEGs across hippocampal cell types (FDR-adjusted P < 0.05, log2 fold change > 0.1), each cell type within each biological replicate was downsampled to an equal number of nuclei (588 nuclei per cell type per mouse). For DG granule cell DEG analyses, each biological replicate was downsampled to ensure equal number of cells per condition (3,041 mature granule cell nuclei per mouse; three mice per condition).

Gene ontology analysis was conducted using Metascape^87^ with genes upregulated by 3 weeks of fluoxetine (expressed in ≥5% of cells; adjusted P<0.05) used as input and all genes detected in DG granule cells used as background genes. Transcription factor enrichment analysis was conducted using ChEA3^88^ with genes upregulated by 3 weeks of fluoxetine (expressed in ≥5% of cells; adjusted P<0.05) used as input.

#### Developmental Plasticity and Regeneration Associated Gene scores

Module scores were generated using the AddModuleScore function in Seurat as described by *Tirosh et al., 2016*^89^. Briefly, AddModuleScore calculates the average expression of the query gene set for each nuclei and subtracts the aggregated expression of matched control genes randomly sampled from genes with similar expression levels. Module scores were visualized across the granule cell population, including both mature and newborn granule cells. For quantitative comparisons, z-scores were calculated within each biological replicate using only mature granule cells and summarized in bar plots.

To generate the “Developmental Plasticity” module score, granule cell neurons were first identified in a published DG single cell RNA seq dataset (*Hochgerner et al., 2018*)^31^ based on expression of neuronal markers (*Rbfox3*, *Slc17a7*) and granule cell markers (*Prox1*, *Dock10*, and *Stxbp6*) (see also Fig. S3A-B). Differential expression analysis was then performed between granule cells from P5 versus P132 mice, and the top 100 genes upregulated at P5 (expressed in ≥5% of cells; MAST test) were used as input for AddModuleScore in Seurat.

To generate the “Regeneration Associated Gene” module score, we used a gene set from *Chandran et al.,* (*2016*)^39^ by selecting genes in the WGCNA “magenta” module, which was upregulated in regenerating dorsal root ganglion neurons following sciatic nerve injury. This list was then filtered to genes annotated as related to neuronal and axonal regeneration based on the authors’ PubMed literature search (Table S2b from *Chandran et al., 2016*). Genes that were also detected in DG granule cells in our dataset were used as input for AddModuleScore in Seurat (see also Fig. S3C).

#### Analyses of External Single Cell and Bulk RNA-seq Datasets

To quantify a “Developmental Plasticity Score” in mature granule cells following environmental enrichment, we analyzed a published hippocampal single nucleus RNA seq dataset (*Pérez et al., 2024*)^45^ comparing mice housed under environmental enrichment versus standard conditions. Mature granule cells were identified by *Dock10* expression among neuronal nuclei (*Rbfox3*, *Slc17a7*) and by the absence of oligodendrocyte-lineage markers (*Olig2*, *Mbp*, *Pdgfra*) (see also Fig. S6A). The top 3,000 most variable genes were used to recalculate PCs and generate UMAP embeddings. Module scores were calculated in mature granule cells for each replicate (n=1 sample per condition; 3 mice pooled per sample) and summarized in bar plots.

For analysis of external transcriptomic datasets in Fig. S4, previously published single cell and bulk RNA seq datasets were imported into Seurat and subsetted to the relevant cell types and experimental conditions (see **Table S3** for full details of each dataset and experimental conditions). Differential expression analyses were performed using FindMarkers (genes expressed in ≥5% of cells; MAST test) and genes were plotted in R by log_2_ fold change relative to each dataset’s respective control groups.

For analysis of *SOX11* expression in human bulk transcriptomic datasets, we obtained TPM normalized expression data from *Huttner et al., 2014* (GSE56267) using GEO2R and the BrainSpan Atlas of the Developing Human Brain (Allen Brain Institute).

#### Dataset 1 and Dataset 2

The above methods describe isolation and analysis pipelines for Dataset 1. Dataset 2 (Fig. S5) followed the same overall workflow, but with two key differences: 1) four experimental groups were included (vehicle, corticosterone, corticosterone + fluoxetine, and corticosterone + electroconvulsive shock; see also “*Electroconvulsive shock (ECS)*” below), and 2) hippocampal neuronal nuclei were enriched using fluorescence activated nuclei sorting rather than processing bulk hippocampal homogenate (see also “*Flow cytometry*” below).

For Dataset 2, purified neuronal nuclei were profiled using the 10x Genomics Single Cell Gene Expression 3’ platform (v3), following the same general protocol and reagents as in Dataset 1. Nuclei hashing was performed to barcode biological replicates with CellPlex Multiplexing Oligos (CMO; 10x Genomics, 3’ CellPlex Kit Set A, 1000261) according to manufacturer instructions (see also “*Flow cytometry*” below). Barcoded nuclei were pooled, and ∼15,000 nuclei were loaded into each of four wells of a Chromium Chip A for droplet generation with barcoded gel beads using the Chromium Controller. Libraries were prepared by the Single Cell Analysis Core of the Columbia University Sulzberger Genome Center following manufacturer instructions and sequenced on an Illumina Novaseq 6000. Libraries were sequenced to a mean depth of 45,000-60,000 reads per nucleus with ≥33% sequencing saturation. Reads were processed using Cell Ranger v6.1.2 and aligned to the mm10-2020-A mouse reference genome. Approximately 70% of all nuclei were successfully assigned to a CMO barcode using the “cellranger multi” pipeline, resulting in 25,974 recovered nuclei for downstream quality control processing.

Downstream quality control, clustering, and gene expression analyses for Dataset 2 were performed in R v4.3.0 using Seurat (v4.4.0), following the same analysis pipeline as Dataset 1. Doublets were defined as nuclei assigned to more than one CMO barcode (∼5% of nuclei) and were removed prior to downstream analyses. Nuclei with <1,600 or >3,600 detected genes, or >5% mitochondrial reads, were excluded. Genes detected in fewer than five nuclei were also removed. After filtering for these quality control metrics, 22,199 nuclei were retained (5,919 vehicle; 6,313 corticosterone; 5,323 corticosterone + fluoxetine; and 4,644 corticosterone + ECS; see also Fig. S5).

#### Data Availability

Analysis scripts can be found on Github (https://github.com/pnguyen1003/snRNA-seq-analyses). Original data for Datasets 1 and 2 have been deposited in GEO under ascension numbers GSE289328 and GSE289329, respectively.

### Electroconvulsive Shock (ECS)

Mice were anesthetized with 2% isoflurane using a nose cone with oxygen flow rate of 1 L/min. Electroshocks were delivered to anesthetized mice through ear clipped electrodes at 120 Hz and 50 mA, for a duration of 1 second. Ringers salt solution was applied to the ear prior to shock to improve conductance. Mice received electroconvulsive shock (ECS) treatment once every other day for a total of 10 sessions and euthanized a day later (21 days) or received 1 shock and euthanized a day layer (1 day). For single nuclei RNA-seq experiments in Dataset 2, all groups (vehicle, corticosterone, cort + fluoxetine, cort + ECS) were treated in parallel such that library preparation and sequencing were performed in one batch. The vehicle group received drinking water with 2 mg/mL beta-cyclodextrin (see “*Pharmacological approach*”).

### Flow Cytometry

For fluorescence activated nuclei sorting (FANS), flow cytometry was performed as previously described^74^. Briefly, mice were perfused transcardially with ice-cold 1x PBS,and intact hippocampi were dissected under a dissecting microscope with meninges and excess myelin removed. Hippocampi (two per mouse) were mechanically homogenized on ice in 1% BSA-PBS (Sigma-Aldrich, A3059) using a 2 mL glass tissue homogenizer (Kontes Glassware) with 12 strokes of the tight pestle. The resulting suspension was filtered through a 40um cell strainer and nuclei were pelleted at 300g for 5 min at 4°C. Nuclei were resuspended in 22% Percoll (GE Healthcare) in PBS and centrifuged at 900g for 20 min (acceleration 4; deceleration 1) to remove debris. Pelleted nuclei were washed in PBS, blocked in 1% BSA-PBS for 5 min, and stained on ice for 10 min with mouse anti-NeuN-Alexa Fluor 488 antibody (EMD Millipore, MAB377X; 1:1,000). Nuclei hashing was performed using CellPlex Multiplexing Oligos (CMO; 10x Genomics, 3′ CellPlex Kit Set A, 1000261). CMOs were added following primary antibody incubation and samples were incubated for an additional 5 min at room temperature. After two additional PBS washes, biological replicates were pooled prior to fluorescence activated nuclei sorting. All buffers contained Protector RNase Inhibitor (Sigma-Aldrich, 03335399001) at 0.2 U/µL for downstream single nucleus RNA sequencing. DAPI (Sigma-Aldrich, D9542) was added to each sample immediately prior to sorting to identify nuclei. Nuclei were sorted on a Sony MA900 cell sorter and gated by forward/side scatter, DAPI, and NeuN (see Fig. S5D for gating strategy). Flow cytometry and sorting were performed at the Columbia Stem Cell Initiative Flow Cytometry core facility at Columbia University Irving Medical Center under the leadership of Dr. Michael Kissner. All data analysis was performed using FlowJo software.

### Immunohistochemistry and Fluorescence in situ Hybridization (FISH)

For immunohistochemistry experiments, mice were perfused transcardially with ∼10 mL of ice-cold 1X PBS followed by ∼10 mL of 4% (weight/volume) paraformaldehyde in PBS. Brains were post-fixed in 4% PFA for at least 4 hours and then cryoprotected in 30% sucrose for a minimum of 2 days. Brains were frozen on a BFS-40MPA Freezing Stage (Physitemp) and sectioned coronally at 40 um using an HM450 microtome (Epredia).

Free floating brain sections were blocked in 10% normal goat or donkey serum (Thermo Fisher) with 0.4% TritonX (Sigma-Aldrich) in 1X PBS. Primary antibodies were diluted in 3% normal goat or donkey serum in 0.4% TritonX and incubated with sections overnight at 4°C on a shaker. Secondary antibodies were diluted in 3% normal goat or donkey serum and incubated with sections for 2 hours at room temperature on a shaker. For nuclear labeling, DAPI (Sigma, D9542) was added during the final 5 min of the secondary incubation at a final concentration of 4 µg/mL. Sections were mounted on coverslips with ProLong Glass without DAPI (Thermo Fisher) for super-resolution imaging and ProLong Gold (Thermo Fisher) for all other experiments.

For SOX11 (Sigma, MABE1929, 1:500), BDNF (Proteintech, 28205-1-AP, 1:1000), and PROX1 (R&D Systems, AF2727) immunostaining, antigen retrieval was performed using IHC-Tek Epitope Retrieval Solution (IHC World, IW1100). Brain sections were heated to 95°C for 40 minutes, washed 3x with PBS, and then incubated in blocking solution as described above. Antigen retrieval was not required for WFA (Sigma-Aldrich L1516-2MG; 1:1000), TNR (R&D Systems AF3865; 1:500), DCX (Synaptic Systems 326014; 1:1000), SV2C (Synaptic Systems 119202, 1:500), dsRed (Takara 632496; 1:2000), and ZNT3 (Synaptic Systems 197004; 1:1000). For co-staining with reagents negatively affected by prolonged antigen retrieval (WFA), the antigen retrieval step was shortened to 15 min to optimize signal-to-noise and preserve staining quality for all antibodies. Attempts to detect SOX11 and BDNF were unsuccessful using the following antibodies, both with and without antigen retrieval: SOX11 (Sigma-Aldrich ABN105), SOX11 (Sigma-Aldrich MABE1935), SOX11 (Abcam EPR8192), BDNF (Santa Cruz Biotech sc-65514), BDNF (Invitrogen PA5-85730), and BDNF (Invitrogen PA5-95183). All secondary antibodies were purchased from Invitrogen.

FISH experiments were performed using the RNAscope Multiplex Fluorescent Reagent Kit v2 assay (ACD Bio) according to manufacturer instructions for fixed-frozen tissue, with the exception that sections were not baked for 30 min at 60°C prior to dehydration. Brains were embedded in OCT following 30% sucrose treatment, frozen at −80°C for at least 24 hours, and sectioned coronally at 15 µm on a cryostat. Sections were then mounted on coverslips for RNAscope processing. Mouse *Bdnf* RNAscope Probe (ACD Bio 424821) and TSA Plus Cyanine3 (Perkin Elmer) was used to detect *Bdnf* transcript. For combined RNAscope and immunohistochemistry, sections were immediately blocked and incubated with primary and secondary antibodies (as described above) after completion of HRP-TSA signal development and four subsequent washes.

### Confocal Microscopy and Image Analysis

Histological sections were imaged on a Zeiss LSM 800 laser scanning confocal microscope with Airyscan super-resolution module using a 63x, 40x, or 20x objective. Images were acquired in Zen software (Zeiss) with identical laser power and detector gain settings across experimental groups. All quantifications were performed in ImageJ/Fiji using identical settings applied across images with the experimenter blinded to image identity.

#### Super-resolution Imaging of Mossy Fiber Terminals and ECM Density

For quantification of ZNT3+ and BDNF+ terminal densities and perisynaptic net (WFA+) density, images were acquired in the DG hilus using a 63x objective (1.4 NA) at 1x optical zoom (0.04 um/pixel) in Airyscan super-resolution mode. Quantification was performed in ImageJ/Fiji by thresholding each channel and measuring the area of positive pixels. Thresholds were held constant across all images. Channel density was calculated as percent area (positive area/total area). The total area was defined as the field of view minus the area occupied by large DAPI+ nuclei (putative hilar mossy cells and interneurons). These nuclei were outlined as regions of interest (ROIs) and subtracted using the ROI Manager to obtain the inverse area (i.e. the total area minus the area corresponding to nuclei).

For quantification of mossy fiber terminal morphology, images were acquired in the DG hilus using a 63x objective (1.4 NA) at 1.4x optical zoom (0.04 um/pixel) in Airyscan super-resolution mode. Z-stacks were acquired with a step size equal to half the optical section thickness. Mossy fiber terminal numbers were quantified by measuring at least 20 µm of an axonal branch and manually counting boutons. Bouton area was quantified by outlining each bouton with an ROI and measuring area in ImageJ.

#### Bdnf mRNA Puncta Density

RNAscope puncta were imaged in Airyscan super-resolution mode using the same acquisition settings as for mossy fiber terminal morphology above. Puncta were manually counted using the multipoint tool in ImageJ and normalized to the area of the granule cell layer, defined by PROX1 immunopositive signal.

#### Sox11 Reactivation

Images were acquired using a 40x objective (0.65 NA) at 1x optical zoom (0.08 um/pixel). The proportion of SOX11+ granule cells was determined by manually counting PROX1+ nuclei that were also SOX11 immunopositive.

#### WFA Intensity Across Hippocampal Subregions

For quantification of WFA intensity across hippocampal subfields (in contrast to perisynaptic net super-resolution imaging described above), tiled images were acquired using a 20x objective at 1x optical zoom (0.28 um/pixel) in Zen. Mean WFA fluorescence intensity was quantified in ImageJ by drawing ROIs around each hippocampal subfield and measuring mean intensity. Each value was then normalized to the mean fluorescence intensity of the total hippocampal field of view.

#### Neurogenesis measurements

Tiled images of the DG were acquired using a 20x objective at 1x optical zoom (0.28 um/pixel) in Zen. DCX+ newborn granule cells in the subgranular zone were manually counted using the multipoint tool in ImageJ and normalized to the area of the granule cell layer, defined by DAPI+ signal. For each biological replicate, three hemisections per mouse were quantified and averaged.

### Quantification and Statistical Analysis

GraphPad Prism 10.2.2 was used for most statistical analyses. Statistical tests used are described in text and figure legends. RNA sequencing data was analyzed in R as described above.

